# Protection from epidemics is a driving force for evolution of lifespan setpoints

**DOI:** 10.1101/215202

**Authors:** Peter V. Lidsky, Raul Andino

**Affiliations:** Department of Microbiology and Immunology, University of California, 600 16^th^ Street, GH-S572, UCSF Box 2280, San Francisco, California 94143-2280, USA.

## Abstract

Most living organisms age, as determined by species-specific limits to lifespan^1–6^. The biological driving force for a genetically-defined limit on the lifespan of a given species (herein called “lifespan setpoint”) remains poorly understood. Here we present mathematical models suggesting that an upper limit of individual lifespans protects their cohort population from infection-associated penalties. A shorter lifespan setpoint helps control pathogen spread within a population, prevents the establishment and progression of infections, and accelerates pathogen clearance from the population when compared to populations with long-lived individuals. Strikingly, shorter-living variants efficiently displace longer-living individuals in populations that are exposed to pathogens and exist in spatially structured niches. The beneficial effects of shorter lifespan setpoints are even more evident in the context of zoonotic transmissions, where pathogens undergo adaptation to a new host. We submit that the selective pressure of infectious disease provides an evolutionary driving force to limit individual lifespan setpoints after reproductive maturity to secure its kin’s fitness. Our findings have important public health implications for efforts to extend human’s lifespan.

---

As early as 1889^1^, theories attempted to explain the evolution of the paradoxical phenomenon of aging^2–5^. The majority of theories can be organized in two classes. Non-optimality^2^ theories argue that aging is an inevitable property of life due to somatic^6^ or genetic^7^ damage. Whereas optimality theories propose that aging results from the trade-off between maintenance and reproduction (disposable soma theory)^8^, or that aging is neutral because extrinsic causes of death precede the lifespan setpoints (selection shadow)^7^. Further, aging has also been proposed to result from detrimental side effects of genes that are beneficial in early stages of development (antagonistic pleiotropy)^9^. Additional theories proposing that aging itself is adaptive have not been well supported by either theoretical or experimental evidence^5^.

Taking together these theories are consistent with many lifespan phenomena, however each of them, if taken individually, only explain a subset of observations. For example, the existence of single gene mutations that individually can extend lifespan^10^ is not consistent with non-optimality theories given that both somatic or genetic inevitable damages are unlikely to be controlled by a single “master” gene. On the other hand the current optimality theories cannot explain why the ecological cases that favor the biological immortality are so infrequent, that it is almost never observed. If aging itself is adaptive, and therefore genetically programmed, immortal individuals should be detected due to mutations in genes responsible for aging mechanisms. Another general problem with adaptive aging theories – benefits from individual’s death allocated to its kin should exceed the price of its own life. Some examples of abnormally long-living species (e.g. tuatara, bowhead whale, proteus) can be explained by most existing theories, while some (birds, bats, naked mole rats) – only by few of them (see Extended Data Table 1 for detailed discussion). Attempts to integrate different aging concepts did not yet resulted in a comprehensive unifying theory^4^. Here we propose a kin-selection evolutionary theory, whereby limiting the lifespan-setpoints of a population provides a selective advantage that helps control the impact of infectious diseases and reduces the likelihood of zoonotic transmission where pathogen adaptation to a new host is required to establish epidemics. The theory proposed here is general in the sense that explains the majority of the lifespan-related phenomena (Extended Data Table 1), provides a rational for the absence of immortal organisms, as well as how specific lifespan setpoints are selected to benefit individual kin.

To examine whether lifespan setpoints are linked to protection from infection, we developed a agent-based *in silico* model that incorporates lifespan into a previously developed host-pathogen theoretical framework^11^. In the initial simulation experiments, we considered populations of individuals with a range of defined lifespans. Time units we use are arbitrary and might reflect days, months or years depending on a given organism. All of our conclusions are nicely scalable along the time axis since pathogens tune the length of their life cycles according to the host’s longevity. These were exposed to ten pathogen species that are able to infect susceptible individuals independently of each other (Fig. 1, Extended Data Fig. 1). For simplicity, efficiency of transmission and the fitness penalty produced by each of these ten pathogens were assumed to be identical. The fitness of each infected individual declined gradually up to a 20% maximum penalty for a single pathogen infection. Therefore, an individual’s fitness was reduced to zero if infected with five pathogens (20×5=100), leading to its death. The model assumes that there are no other reasons for death except disease caused by infection or reaching the maximum age span, there is no recovery from diseases, neither vertical pathogen transmission. In addition, an individual’s reproduction efficiency is set to correlate linearly with its fitness, i.e. uninfected individuals have more progeny (Fig. 1a). We also assumed a maximum population number for a single ecological niche of ten thousand individuals. Infection efficiency depends directly of density of individuals in a population, which is deemed to be uniform and ideally mixed using previously described conditions^11^. The model examines the effect of different pathogen transmission rates and assumes that pathogens are never fully eliminated from the system: when the last infected host dies, pathogens are reintroduced from another host species or from the environment. In this manner, our model not only considers the progression of existing infections, but also acquisition of new epidemics. Using this model and starting from an uninfected population, we observe that both the population size and the number of pathogens fluctuate in cycles, until an equilibrium is reached overtime (Fig. 1b, c), consistent with previous models and simulations^11^. The oscillation can be rationalized in terms of infected individuals dying reducing the availability of uninfected hosts. We monitored the number of individuals in the population and the pathogen load, as well as the total population fitness calculated as the sum of the fitness of all its members.

**Fig 1.**
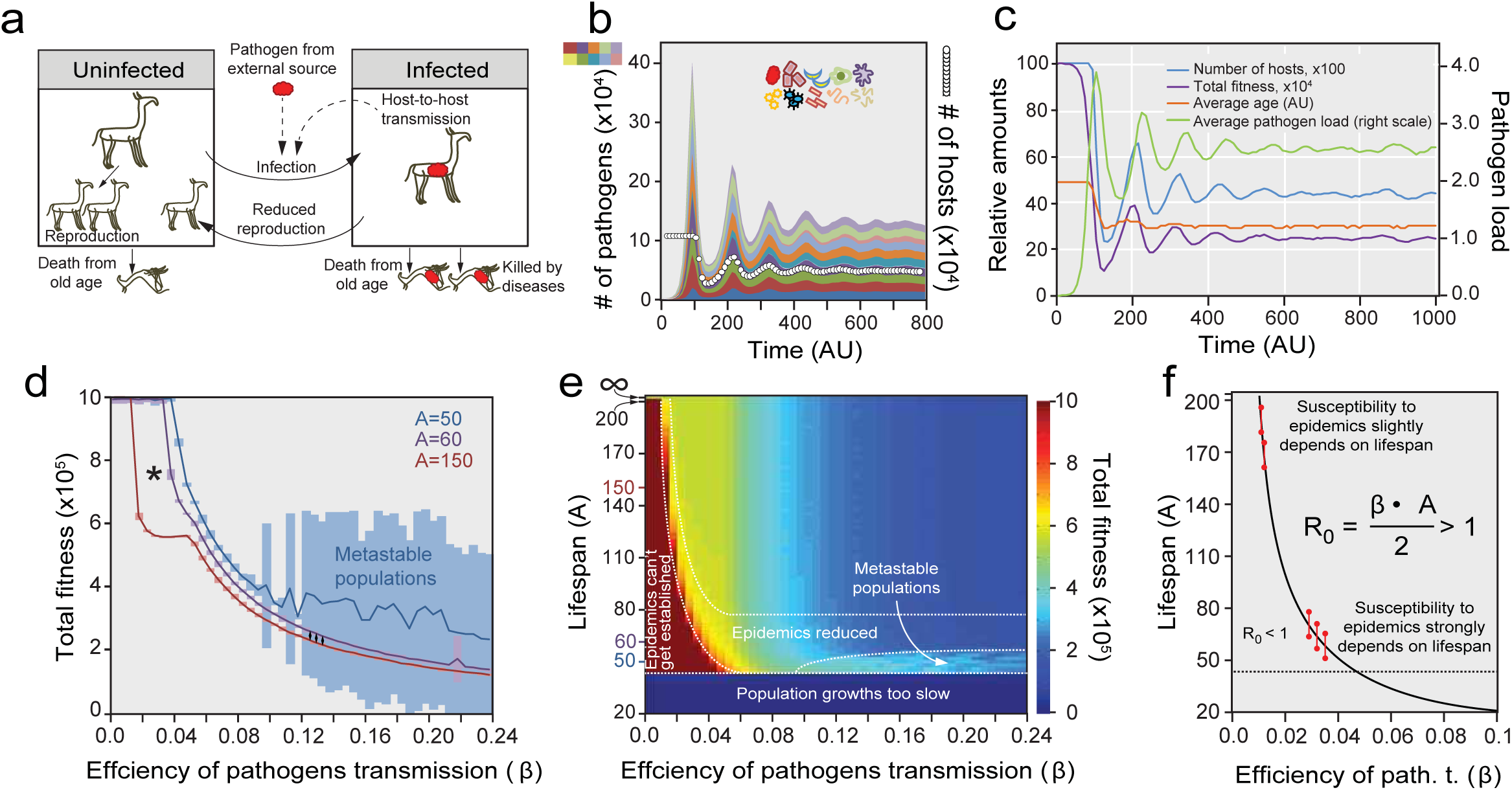
Length of lifespan controls epidemics establishment and progression. **a.** Schematic representation of an epidemic model that considers the effect of lifespan on pathogen transmission. Pathogen transmission is assumed to occur from one individual to another or from the the environment if no infected individuals are left alive. Infected individuals are less efficient in reproduction but give birth to healthy progeny. **b.** Mathematical simulation of pathogen co-circulating in a host population where the lifespan setpoint is 100 time units and transmission efficiency – 0.12. Number of different pathogens is shown. **c.** Results of same simulation as in b. Number of host, average fitness, overall population fitness are represented over time. **d.** Analysis of steady state total population fitness of individuals with different lifespans (A) exposed to pathogens with different transmission efficiencies (β). *A* is the lifespan setpoint of an individual. β - probability of pathogen transmission from an infected to uninfected hosts within a population of susceptible uninfected hosts. The lifespan-dependent ability to establish epidemics is marked with asterisk (*). Region where populations are metastable is seen by extended standard deviation bars. The effects of epidemics reduction due to removal of highly infested elderly individuals is labeled with diamonds (♦♦♦). **e.** Analytic solution of steady state values with different lifespan (*A*) and transmission efficiencies (β). Total fitness of populations is shown by color; other parameters are displayed in Extended Data Fig. 2. **f.** Analytical prediction of the region inaccessible for epidemics establishment. R_0_ – number of individuals in fully packed susceptible population infected by one contaminated host per total time of infection.

We initially analyzed whether different lifespan setpoints impact the total fitness of populations upon infection. We examined pathogens with different transmission efficiencies (β) and individuals with different arbitrary lifespans (A). Total population fitness captured after reaching equilibrium was plotted as a function of lifespan and pathogen transmission efficiency (Fig. 1d, 1e). All lifespan ranges (excluding the biologically immortal control) were indistinguishable with respect to fitness when the pathogen transmission efficiency was very low (Fig. 1e, β = 0.005 to 0.01)). Interestingly, at slightly higher, but still low, pathogen transmission rates (β = 0.015 to 0.045), fitness of the population was negatively correlated with longer lifespans (Fig. 1d, e). Individuals with the longest lifespans render the population more susceptible to poorly transmitted pathogens, compared with a population of individuals with shorter lifespans (Fig. 1d, asterisk). Thus, as individual lifespan shortens, population fitness increases (Fig. 1d, red zone) when faced with low transmission infections. This analysis suggests that longevity decreases population fitness due to the increased susceptibility to epidemics outbreaks (Extended Data Fig. 2c).

**Fig 2.**
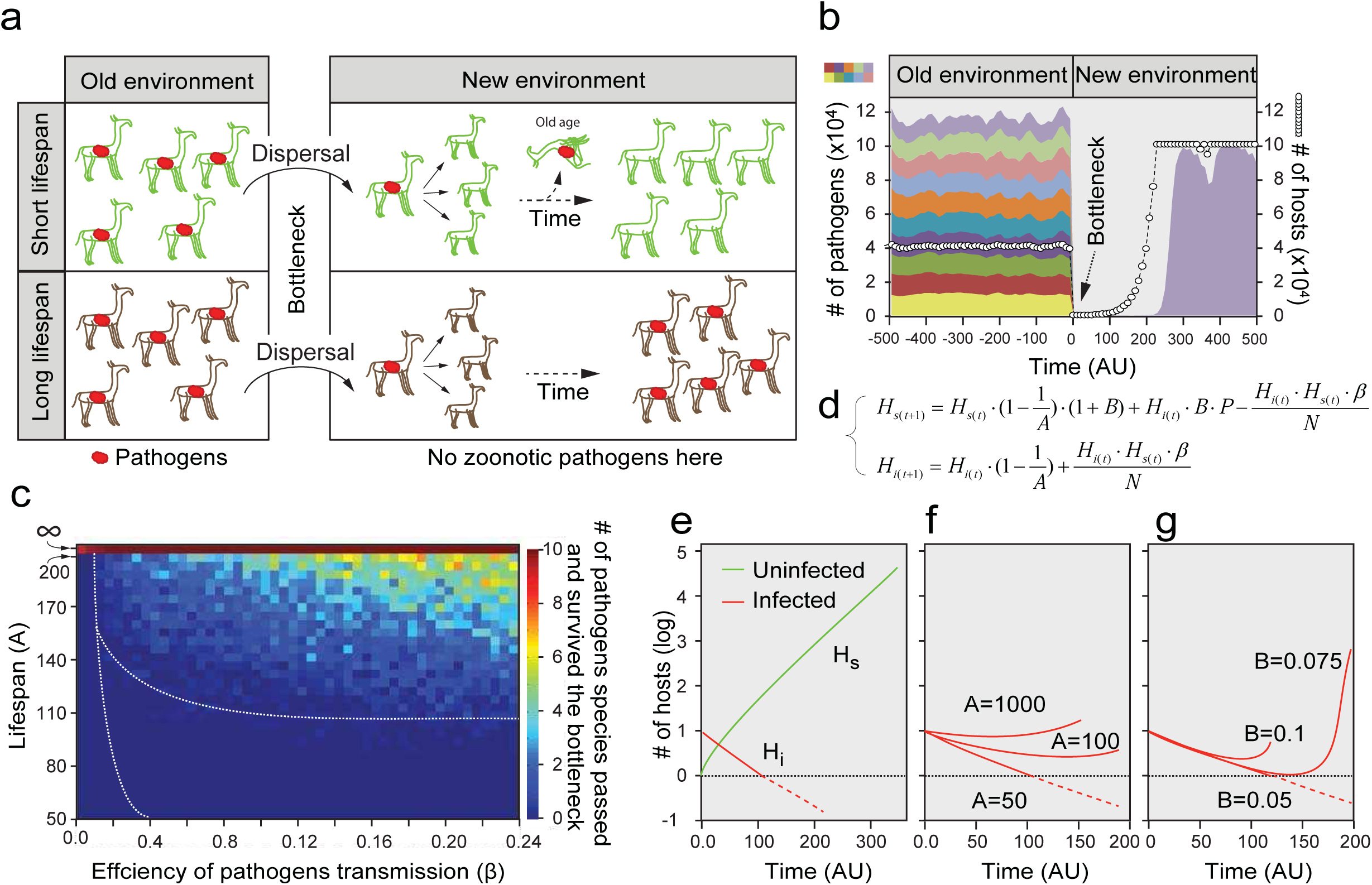
Lifespan limitation facilitate pathogens clearance during bottleneck host dispersal. **a.** Pathogen clearance model following host migration to a new environment. Infected individuals of short lifespan die before population reached the density sufficient for epidemics expansion. As a consequence, pathogens suffer severe bottlenecking and therefore are clear from the population in the new niche. **b.** An example of a single simulation. Random 10 individuals from highly infested population were released into a sterile compartment. Age of programmed death is 150 time arbitrary units (AU), transmission efficiency – 0.13. **c.** Systematic analysis of number of epidemics that survive bottleneck dispersal. Solid line corresponds to one in Fig. 1e, dotted line is a prediction of border age able to clear pathogens, produced from deterministic model shown in panel d**. d.** Analytic solution using a deterministic model. *H*_*s*_ and *H*_*i*_ are numbers of susceptible and infected hosts correspondingly, *B*-birth rate, *P*-pathogen effect on birth rate, *N* – maximum population density, other variables as in Fig. 1d. **e.** Simulation results of equations shown in *d.* If *log(H_i_)* is below zero, we considered pathogen is cleared from the population. (*i)* Initial input parameters were: *A*=50, *B*=0.05, *H*_*s(0)*_=0, *H*_*i(0)*_=10, *P*=0.5, *N*=10^5^, β=0.05. **f.** An increase in age of programmed death results in inability to clear pathogens via bottleneck dispersal. Same experiment as in panel e was performed with different values of *A.* **g.** Effect of fecundity on pathogen clearing. Simulation results with different values of *B* reveals negative correlation between fecundity and clearing efficiency (see also Extended Data Fig. 3h, i for more detailed analysis).

An analytic solution to the model also supports these conclusions. The basic requirement for pathogens to spread in the population is that the number of susceptible individuals infected by pathogens shed by a single infected host (herein R_0_) – should exceed 1. R_0_ can be expressed as transmission efficiency (β) multiplied by duration of the infection, which we initially assumed to average half of the host lifespan 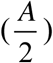. For simplicity, the population density-related component is considered to be 1 and it is omitted from the calculations. The interdependence 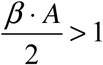 or 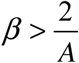 demonstrates that the establishment of infection at low transmission efficiencies is determined by lifespan (Fig. 1f). If the lifespan is shorter than the time required for transmission, pathogens are unable to establish an epidemic. Thus, populations composed of shorter-lived individuals are expected to be more protected from invasion of pathogens, particularly if pathogens have lower transmission rates (β = 0.005 to 0.045). These results suggest that the benefit of a limited lifespan is to protect populations from low or moderately transmissible diseases that establish persistent infections; for humans many such persistent infections are common, including AIDS, hepatitis B and C, leprosy, herpes, tuberculosis, and helminths.

Even if an epidemic is established, i.e. when pathogen transmission efficiency becomes high (β = 0.1 to 0.24), populations with shorter-lived individuals also experience a fitness benefit arising from the *metastability* of host and pathogen populations (Fig. 1d, e, light blue zones highlighted as “metastable populations”). The benefit results from the significant reduction in host population density caused by rapid spread of the infection, which, when combined with shorter lifespans, leads to a dramatic decline in pathogen numbers thereby allowing the population to recover from infection (Extended Data Fig. 3e-f).

**Fig 3.**
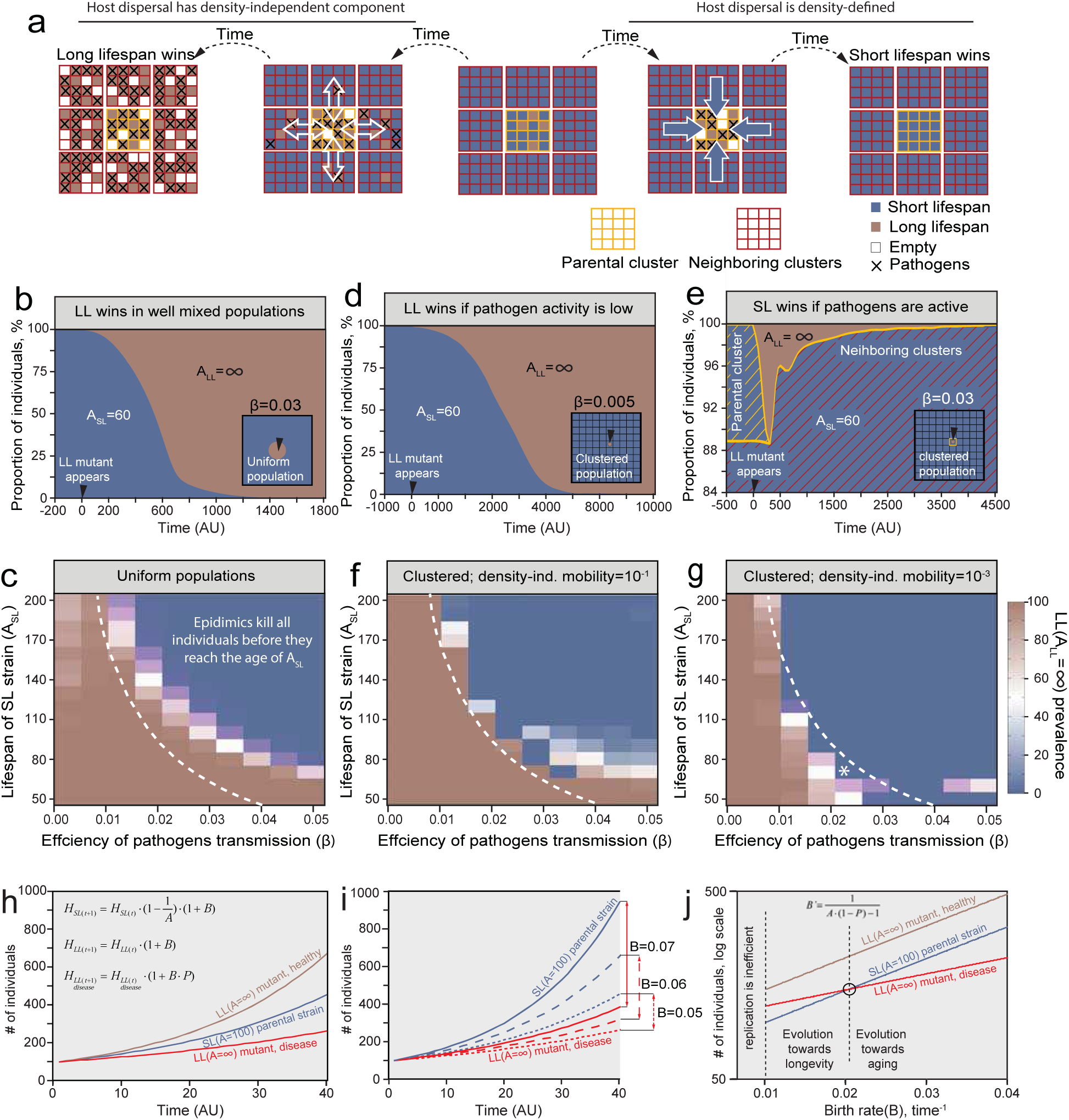
Asymmetric dispersal. **a.** Schematic representation of the host asymmetric dispersal or migration. **b** and **c.** Long-living (LL) individuals displace short-living (SL) individuals in uniform population. **b.** A small group (10) of long-lived (A=∞) individuals is introduced at time 0 into a population of short-lived (A=60) individuals. **c.** Systematic analysis reveals that region where shorter lifespan gives benefits (below the dashed line, taken from Fig. 1e) is susceptible to invasion of long-living mutants in uniform populations. Percent of long-living individuals is shown. **d** to **g.** Analysis of clustered metapopulations. **d.** Long-living strain invade efficiently if pathogens transmission rate is low (β=0.005). **e.** In presence of higher pathogens transmission rate (β=0.03) shorter-living strain displace the longer-living individuals. Parental cluster (yellow hatched area) and new clusters (red hatched area). Note individuals from the parental cluster are eliminated and mutants arise (yellow hatched area). **f** and **g**. Systematic analysis of competitions in clustered metapopulations reveals a region (* in **g**) where the long-living mutant can not invade and short lifespan is beneficial. The existence of this region strongly depends on low levels of density-independent dispersal. Percent of individual simulations where long-lifespan mutant took over is plotted. **h-j**. Individual’s birth rate strongly affects asymmetry between short-living and long-living infected populations densities. **h.** Deterministic equations describing population growth of short-living, long-living and long-living infected individuals. Variables are used exactly as in Fig. 2d. Simulation of population growth. *A*=100, *H*_*0*_=100, *B*=0.05, *P*=0.5. **i.** Higher birth rates are resulting in increased spread between short-living and long-living infected population growth. Three pairs of curves are shown. **j.** Lifespan prediction. Simulations essentially as in **j** were run with different values of *B*. Results at *time*=40 are shown. At small values of *B* populations of infected long-living individuals are replicating faster than populations of healthy aging individuals, therefore favoring fixation of longer lifespan. At larger values of *B* healthy short-lived individuals can overgrow infected long-living ones, thus favoring the former strain. Equilibrium point, where two lines were crossing (black circle), was found analytically at 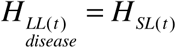 from the equations shown in *h*. Therefore in this simplified condition the equilibrium lifespan A’ can be estimated from known *B* and *P* (equation).

Populations often migrate to colonize new environments. To further study the role of lifespan setpoints in controlling infection, we next examined the relationship between lifespan and pathogen load in the case of host populations dispersing to a new environment. In our modified simulation and analytical models, dispersal involves extreme population size reduction, “bottlenecking”^12^, whereby a small group (10) of random individuals from infected population colonize the new environment (Fig. 2a, b). Under these conditions, since the population density is low, pathogens will not spread efficiently soon after migration. The model predicts that in populations with shorter lifespans, the infected founders will die before the population density has reached a level that allows for efficient transmission. Consequently, populations composed by shorter lifespan individuals more efficiently clear pathogen than populations composed of longer-lived individuals (Fig. 2b-f). Of note, clearance efficiency also depends on the basal fecundity rate (*B*-number of progeny produced by 100% fit individual per time unit). Populations with highly fecund individuals will reach high densities faster and, thus, the probability of pathogen spread in a population by infected founders will increase accordingly (Fig. 2g, Extended Data Fig. 3h, i).

We also observed a minor positive effect of lifespan limitation on the fitness of highly infected populations as shown previously^13,14^. Since older animals possess more chronic pathogens than younger ones, their removal could increase the total population fitness (Fig. 1d, diamonds, 1e, “epidemics reduced”; Extended Data Fig. 3a-d). Importantly, our model suggests that this is a minor effect, compared to the more significant consequences resulting from epidemics prevention and pathogen clearance following host population bottleneck dispersal (Fig. 1d-f, and Fig. 2).

Individuals that live longer should reproduce for longer times and be more effective in passing on their genes. Thus, the evolution of traits that limit lifespan, common to most organisms, is counter-intuitive. We thus examined the whether long-living variants can displace a population of short-living individuals. To this end, we modified our simulation to allow for variants with a very long lifespan (A=∞) to emerge in a population of individuals with a defined lifespan (e.g. A=60). Under conditions where all individuals in a population intermix evenly, and thus long and short lifespan individuals have equal probabilities to interact with each other, the longer-living variants efficiently outcompete shorter-living individuals (Fig. 3b), even if this leads to establishment of epidemics (Extended Data Fig. 4a). However, under conditions of severe epidemics, premature death caused by pathogens equilibrated the number of short and long-lifespan individuals, thus limiting the advantage of long-lived variants (Fig. 3c, blue zone, Extended Data Fig. 4b, c). Populations located in the region of parametric space with short lifespan-associated benefits, i.e. the region where epidemics can’t get established and metastable populations (Fig. 1e), were susceptible to displacement by longer-living individuals (Fig. 3c, Extended Data Fig. 4b, c, area below the dashed line). Thus, in ideally mixed populations long-living variants will displace short-living individuals.

**Fig 4.**
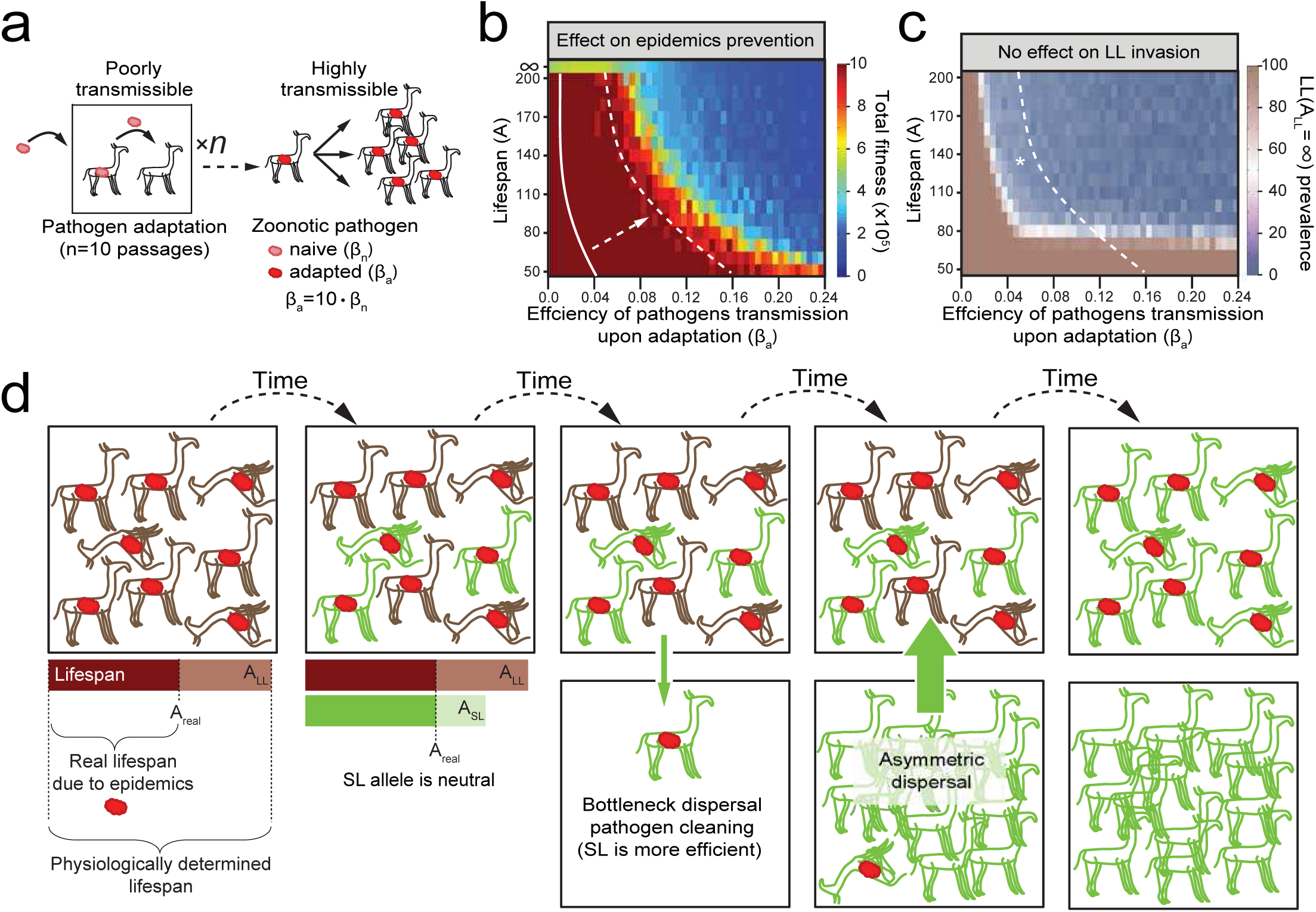
The role of pathogen adaptation in lifespan selection. **a.** Schematic representation of pathogen adaptation in the context of zoonotic transmission. **b.** Pathogens misadaptation results in significant increase of the area on parametric space where shorter lifespan is beneficial. Experiment was performed as in Fig. 1d, but initial pathogen transmission was decreased 10-fold relatively to values shown on the axis. Pathogens required 10 passages to reach its adapted state and increase transmission. Dashed line is from Fig. 1f. **c.** Pathogen misadaptation does not affect invasiveness of long-living mutant. Asterisk labels the continuous area where non-aging mutant cannot invade (blue) and aging gives selective advantages (below the dashed lines) at the same time. **d.** Evolution of shorter lifespan or origins of aging. Fixation of shorter lifespan demands an ongoing epidemics as a prerequisite. Under this conditions replication success of short-living individuals is equal to those of their long-living counterparts. However short-living animals have a selective advantage for cleaning pathogens via the bottleneck dispersal. Healthy short-living individuals might displace long-living strain from the parental cluster due to asymmetric dispersal process.

The conclusions are different when populations are composed of sub-populations or clusters that are spatially separated, for instance by living in different niches. These spatially separated sub-populations may interact by exchanging individual members that move from one sub-population to another. We studied competition between longer-lived and shorter-lived organisms in this context of clustered populations, by modifying our model and introducing barriers between subpopulations (Fig. 3, Extended Data Fig 5). We based our analysis on a previously proposed computational model used to explain the seemingly altruistic effect of programmed cell death in unicellular organisms^15^. We observe that in the context of infection of clustered populations, the shorter-lived individuals outcompete long-lived variants. As a result of infection, the density of individuals in uninfected clusters is higher than in infected clusters. Accordingly, uninfected individuals from overcrowded clusters are more likely to migrate to infected clusters where the density is lower. This “asymmetric dispersal” model proposes that the direction of dispersal indirectly depends on the fitness of the individuals in a given cluster, which is linked to infection (Fig. 3a).

Using this model, we initially performed a source-sink density-dependent dispersal simulation^16,17^ consisting of a 10 x10 grid of niches with capacities for three hundreds individuals each (Extended Data Fig. 5b). We introduced additional fitness penalties on reproduction to simulate the overpopulation effects and density pressures more adequately (Extended Data Fig. 5a for details). As mentioned, asymmetry in population densities between clusters forces migration of individuals from high density niches to a less populated cluster. At the start of the experiment all clusters were populated with individuals of a given limited longevity. Then a small group of long-lived (A=∞) mutants was introduced into one of the central clusters (Fig. 3d, e, Extended Data Fig. 5b) and the competition between genotypes was simulated. Our analysis revealed that when pathogen activity is low (β=0.005) (Fig. 3d) long-lived mutants displace the short-lived individuals with efficiencies comparable to those in uniform populations (Fig. 3b). If we examine this process considering density-independent dispersal, with individuals migrating at a high frequency (e.g. birds or bats, Fig. 3f, mobility 10^−1^), we find that clusters behave as a well mixed environment (similar to Fig. 3b-d). Thus, longer lifespan will also be selected in populations of highly mobile individuals (Fig. 3f, Extended Data Mov. 1). In contrast, when we consider a density-dependent dispersal under higher pathogen transmissibility conditions (β=0.03), shorter lifespan individuals displace long-lived variants (Fig. 3e, Extended Data Mov. 1). Our analysis further revealed an area where the short lifespan is beneficial and long-living strains are not able to prevail (Fig. 3g, *). Our model also suggest that upon infection, the emergence of longer-lived mutants also leads to extinction of the shorter-lived individuals in that cluster (Fig. 3e, dashed with yellow); as a result, the generation of longer lived mutants is detrimental to all individuals within the cluster.

We next examined the role of fertility. Asymmetry in population numbers between healthy short-lived and infected long-lived population strongly depends on their fecundity (Fig. 3h-j). Thus, using deterministic equations (Fig 3h), we modeled growth of short-lived and longer-lived populations with arbitrarily distinct birth rates (*B*). We propose that short-lived individuals outcompete long-lived ones in a birth rate-dependent manner, because high fertility allows asymmetry in density between clusters to develop more rapidly (Fig. 3i). We further propose the existence of an equilibrium when the population growth of healthy short-lived individuals approach that of infected long-lived individuals (Fig 3j, black circle). Our model predicts a simple interdependence between fecundity, lifespan and pathogen effects on fitness (*P*) (Fig. 3j, equation). This dependency, together with fecundity effects during bottleneck dispersal (Fig. 2g, Extended Data Fig. 3h, i) explains the negative correlation between longevity and fecundity, as previously proposed in the context of the disposable soma theory^8^.

An important source of new pathogens in natural populations is the horizontal transmission between species, also called zoonotic transmission. The species-to-species barrier often requires that pathogens undergo several cycles of replication in the new hosts before adapting and gaining optimal transmissibility (Fig. 4a). We next considered how longevity impacts the ability of zoonotic infections to become established in a population, by modeling how populations with short-lived or long lived individuals influence pathogen adaptation to the host. We used a model similar to that used in Fig. 1e, with the exception that the pathogens are initially attenuated with a 10-fold transmissibility reduction; adaptation requires the pathogen to be passaged between ten host individuals to reach higher transmission efficiency. Under these “zoonotic infection” conditions, we observed that the range of efficiency of pathogen transmission in which epidemics cannot get established is increased dramatically for populations with shorter lifespan individuals (Fig 4b zone below the dashed line), while the invasion of populations with long-living mutants (A=∞) remains unaffected (Fig. 4c). This suggests that the advantage of short lifespan is particularly important in the context of zoonotic epidemics. Thus, our model suggests that asymmetric dispersal and pathogen adaptation to a new host are two critical factors enhancing the benefits and evolutionary stability of shorter lifespan.

Taken together, our analyses indicate that host–pathogen coexistence and coevolution played a key role in determining the lifespan setpoints of a species. A population of longer-lived individuals is more susceptible to introduction of pathogens from other species, less effective to clear the infection following bottlenecking, and is less fit than a population of shorter-lived individuals when epidemics progress. Extension of the lifespan setpoint is beneficial only in the absence of pathogens, which for most species is not a realistic scenario in an evolutionary context. However, tolerance to pathogens (e.g. in bats) might alleviate selection pressure resulting in expansion of lifespan. The asymmetric dispersal from uninfected highly populated niches to infected less populated ones disfavors the emergence of longer lived individuals (Fig. 3). A series of asymmetric dispersal events should then lead to fixation of lifespan-setpoints optimized for the elimination of pathogens from a given population. Importantly, during epidemic outbreaks following asymmetric dispersal, we find that longer-lived individuals place even shorter-living individuals in the same cluster under the risk of extinction (Fig. 3e). Thus, infection is likely to be a major evolutionary force dictating the puzzling near-absence of immortal or extremely long-living individuals in nature. Indeed, subpopulations able to produce such long-lived variants are exposed to the catastrophic risk of being decimated during outbreaks. We thus conclude that lifespan of a species is a result of a trade-off between the pressure produced by pathogens and the pressure towards life extension and fecundity.

It is now generally accepted that host-pathogen interactions constitute major driving force during evolution^18^. The limitation of lifespan, leading to aging, may be one of the earliest and more robust population-level defense mechanisms against the spread of new pathogens and clearance of those that have been already established. A crucial role in the evolution of lifespan may be played by chronic low-transmissible diseases that persist for a significant part of a host’s life and cannot be cleared by immune system. A major selective pressure towards shorter lifespans may also arise from prospective pathogens that are present in the environment but did not yet adapted to a given hosts or established epidemics.

Our work does not negate but complement and extend previous evolutionary theories of aging. For example, our model is consistent with selection shadow and antagonistic pleiotropy theories (Fig 4d, Extended Data Table 1). However, we identify host-pathogen interactions as a major selective pressure driving evolution of lifespan setpoints. It should be also emphasized that our study does not invoke group selection^19^, but rather proposes the selection of shorter lifespan depends on the inclusive fitness of the individuals in the context of the population. Hence we consider evolution of lifespan a kin selection strategy that favors the reproductive success of an organism's relatives, even at a cost to the organism's own survival. Finally, considering lifespan termination as an adaptive trait, our theory does not provide mechanistic insights of aging. The lifespan setpoints likely exploit similar mechanisms across different species, such as the modulation of DNA and protein damage responses, stress responses and senescence pathways^6,20–22^.

Research on aging, searching for lifespan determinants may lead to an effective increase in human and animal lifespans. Our ability to describe and model aging as an evolutionary process linked to infection, provides a new paradigm to identify lifespan mechanistic programs which can be prevented or even reverted. On the other hand, our theory also alerts to potential epidemiological risks associated with extended lifespan (Extended Data Fig. 6h). Conversely development of efficient mechanisms to limit or tolerate infections weakens the evolutional pressure towards lifespan shortening. We propose this as an explanation for abnormal longevity in bats, mole rats (Extended Data Tables 1) and humans.

## Methods

### Data collection and presentation

All simulation scripts were coded and compiled with Python 2.7 (Python Software Foundation). Data was plotted with Excel (Microsoft) and MatLab (Mathworks).

### Models

While our infectious catastrophe theory is well supported by previously published empirical evidence (see Extended Data Table 1a), this manuscript does not include any original experimental data supporting its predictions. Therefore it is crucial to demonstrate that our models are adequate and the values of parameters we use are realistic.

Our models are based on classical host-pathogen interaction models^1^, adapted to agent-based simulations to better account for the individual life-history effects on how epidemics proceed. The basic algorithm is presented in Extended Data Fig. 1. The values and main processes are listed in Extended Data Table 2a, b.

The host’s population sizes we used (10 thousand individuals in uniform simulations and 300×100=30000 in clustered metapopulations) were big enough to exclude any effect of stochastic factors on our final results.

In the beginning of simulations the niches were filled with maximum number of individuals with randomly generated ages (*a*) and replication values (*r*). Generation of uniformly distributed random numbers in all cases was performed with *random.py* Python library (Python Software Foundation) which uses standard Mersenne Twister algorithm.

Host’s reproduction was simulated the following way. Each round a value has been added to the replication value in each host:

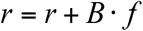

where B is birth rate (0.05 in all simulations, 0.075 and 0.1 in Extended Data Fig. 3h, i) and *f*-individual’s fitness (*f*_max_=100 for an uninfected animal). When *r* was reaching value of *r*_max_=100, the individual gave birth to another genetically identical individual. *r* of both parent and daughter individuals were set to zero. One generation of healthy individuals in typical simulation could therefore be calculated as:

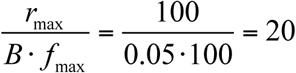 arbitrary time units. New born animal was always considered to be uninfected. If population has reached its maximum (N_max_) in simulations of uniform populations, the newborns were considered to be aborted or expelled, while the *r* of the parental animal was still set to zero. For metapopulation experiments we used a fitness-dependent algorithm of population limits. To increase the pathogen effect on birth rate and to simulate overcrowding effects more adequately, the progeny was delivered to an individual only if 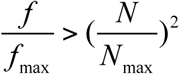. We assume that under conditions of resource shortage only the fittest individuals could leave progeny. However results similar to ours could be obtained with increase of pathogen’s adverse effect on birth rate without changing its effects on lifespan (unpublished).

Dying due to reaching the lifespan setpoint was simulated by incrementing the age of individuals (*a*) each round and comparing it to the preset age of death (A). For simplicity we assumed no visible fitness declines were preceding the individual’s death: very first “symptoms” of aging were considered to be lethal.

As mentioned in the main text, time units we use are arbitrary and might reflect days, months or years depending on a given organism. All of our conclusions are nicely scalable along the time axis since pathogens tune the length of their life cycles according to the host’s longevity.

In this study we considered only chronic pathogens that cannot be cleared by the immune system. Such diseases are numerous and well known. We suggest long-lasting diseases, but not ones with rapid recovery, are contributing to the selection pressure towards lifespan shortening. Simulations in Extended Data Fig. 7d shows that introduction of recovery degrades the benefit produced by lifespan setpoint.

Infections were simulated in the following way: in the first step the infection load of the population (L_pi_) has been estimated for each pathogen as a number of individuals infected with this pathogen. If this value was declining below 1, we assumed the presence of zoonotic pathogen reservoir and set as L_pi_=1. Next the infection efficiency was estimated as β•L_pi_, where β is transmission efficiency analyzed in a broad range of values of 0.005-0.24. To calculate number of pathogen attacks β•L_pi_ has been transformed into integer using stochastic rounding algorithm (e.g. 4.89 was giving 5 in 89% of cases and 4 in 11% of cases). Then these attacks were then randomly firing in the population. If the attack was hitting the susceptible individual it was infected, however if it hit an empty slot or an individual already infected with this pathogen, the attack was considered to have no consequences. Therefore numerically number of newly infected individuals in each round was equal to classical models: β⋅L_pi_⋅H_pi=0_, where H_pi=0_ is a number of susceptible animals in population. The non-canonical algorithm of infection was used to simulate multiple independent infections and pathogen’s evolution (Fig. 4) more easily.

We have used multiple (10) pathogen species in all our simulations. In each experiment all of them had the same parameters. We assumed no interactions between pathogens; so they were able to infect hosts in any combinations. It was important to show that selective pressure towards shorter lifespan could be produced by the cumulative effect of several different pathogens with relatively mild pathogenesis. Multiplicity of pathogens also allowed us to minimize a number of disease parameters and still simulate fertility decline, long chronic infections as well as disease-inflicted death with a true stochastic component. It was also technically convenient, since it reduced noise in our simulations. Usage of a single pathogen with very strong adverse effects (e.g. developing its fitness penalties non-linearly over time, similarly to HIV, hepatitis or syphilis) would immediately raise questions about the universality of the simulation results. Using the parameters of humans diseases listed above would also require introducing vertical transmission as well as many additional conditions and parameters that would further complicate the model. Nevertheless all our conclusions are also clearly valid for such a single “terminator superpathogen” (unpublished). It should be also noted that vertical pathogen transmission (infection of the progeny by the parent immediately upon birth) will strongly increase key effects observed in Figs. 1 and 3 (unpublished).

A penalty that limits a host’s fertility due to disease (*P*) is an accepted fact: resources to be invested in reproduction are hijacked by pathogens and/or reallocated into deployment of immune responses. For analytical experiments (Fig. 2e to g, 3 h to j) we have used a 50% penalty, while in simulations a 20% (elsewhere stated) decrease in fitness per pathogen was used. The latter, more moderate number was selected so as to allow for several pathogens in the system. We did not introduced lifespan shortening by the single pathogen as it would complicate the interpretation in regions with low β. The gradual development of disease severity within 15 time points should be considered as a minor factor, introduced to eliminate some unaesthetic oscillations. In Fig. 1f we assume population density-dependent component to be equal to 1, and therefore 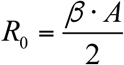. We consider this as an adequate simplification in the region of parametric space, where *R*_0_ ≈ 1. Adverse effects of epidemics are minimal in this region and proportion of susceptible animals in the population is ≈ 1. If the ages in population are distributed evenly, average age of infecting, as well as its duartion is 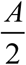. It should be noted that tiny fading outbreaks are happening in stochastic models even if *R*_0_ < 1 (not shown).

In Fig. 2 we demonstrate how infection can be cleaned following the bottlenecking of the host populations. To simulate highly infected starting populations, we have run the long-living individuals (A=∞) until they have reached the steady state adapting all 10 pathogen species. Then 10 random individuals were selected from this population and their lifespan setpoints were changed to the indicated value. Therefore the initial sampling was not affecting the pathogen composition immediately after the bottleneck. It was made from analogous moieties and usually retained all 10 diseases. For simplicity in the main text we discuss the dispersal of small population of founders to a new niche (Fig. 2), free from external pathogens. However taking in account the concept of pathogen adaptation (Fig. 4) it is clear that efficient cleaning from pathogens might follow the population decreases caused by diseases or predator-prey interactions occurring on the same territories: individuals might clean up the well-adapted pathogens, while the exposure to external non-adapted ones will not result in immediate restart of epidemics.

Simulation of competition between long-lived and short-lived individuals was performed in two ways that gave similar results (Extended Data Fig. 4). First, *in situ* generation of long-living mutant was simulated. Populations of short-living animals were run until reaching the steady state. Next, 25 random animals were transformed into long-living mutants (A=∞) and simulations have been running until the new steady state was reached and data was collected. Second, extrinsic invasion of long-living mutants was studied. Two isolated populations of short-living and long-living (A=∞) mutants were run in parallel under identical conditions. Upon reaching the steady-state, 25 slots were randomly swapped between two compartments with period of 100 rounds.

Source-sink density-dependent dispersal simulation (asymmetric dispersal) is one of the key results of the paper (Fig. 3, Extended Data Movie 1). Both empirical and theoretical studies show that dispersal as well as the underlying movement behavior are condition-dependent and informed processes. Local population density is considered to be a major factor affecting dispersal^2–5^. It should be emphasized that activation of a specific program for dispersal upon overcrowding was reported even in bacteria (swarming and quorum sensing)^6,7^ leaving no doubt that our assumption for universality of density-dependent dispersal is valid. Intraspecific competition and resource shortages push excess individuals into neighboring, less populated regions. We also note that purging the long-living mutants occurs rapidly in the course of epidemics progression. We suggest this to be much faster than most of evolutionary processes, since it does not require rare events (mutations) to happen. Taking this into account, we consider that the minimal stringency of physical barriers that separate different parts of population is less than in most other evolutionary models. Therefore our simulations (Fig 3d to g, Extended Data Movie 1) are based on adequate presuppositions. Simulations for clustered metapopulations (Extended Data Fig. 5) were performed first for each cluster as it would be a uniform population, and after density-dependent and density-independent dispersal were simulated. To exclude potential artifacts coming from the same order of cluster analysis, the sequence of cluster calculations was randomized at each round.

Equation if Fig. 3j was obtained from: 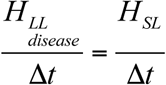. Under these conditions (and if density-independent dispersal is insignificant) the evolution of lifespan does not go neither towards lifespan elongation, nor shortening since the equilibrium between populations is reached. Thus from equations shown in Fig. 3h we obtain: 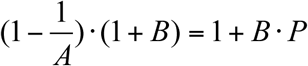, and the birth rate in the equilibrium point: 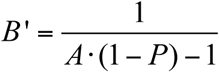. Decrease in birth rate will result in evolution towards longer lifespan and will move B’ to the left, while an increase in birth rate will lead to lifespan shortening and increase B’.

The concept of pathogen adaptation to new species is widely accepted. We can illustrate this with an example of HIV-2. Closely related viruses are persisting in sooty mangabeys (*Cercocebus atys atys*). While most serotypes of HIV-2 are known from single human hosts only, some can also be transmitted from human to human^8^. Thus it seems while most HIV-2 variants that infect humans result in a dead-end, a few are taking an opportunity to adapt enough to establish epidemics. Since HIV pathogenesis occurs in prolonged periods of time comparable to human’s lifespan, we suggest that longevity could assist pathogen adaptation in the end resulting in epidemic outbreaks. To simulate pathogen adaptation (Fig. 4b, c), each pathogen in each host obtained an additional parameter – passage number (x). For pathogens coming from the environment x was set as zero. At the stage of infecting the program assembled an array of passage values from a pathogen’s populations. Representation of each passage in the array was calculated as the number of individuals infected with the pathogen at the given passaging age multiplied by the relative transmission efficiency of this pathogen (β_n_ or β_a_). Next for every successful pathogen attack, the algorithm randomly selected a passage number from this array and incremented it with 1 to assign to a newly infected host. Therefore the composition of a pathogen’s metapopulation was carefully simulated.

Simulation data was collected upon the epidemics was reaching steady-state. In experiments with uniform populations 3×10^3^ of rounds were sufficient. For simulations of lifespan variants competition in uniform populations we used 9×10^3^ rounds after mutant introduction. In clustered metapopulations the run was stopped after one of the variants was winning.

Number of replicate simulations is at least 10 per data point in all heatmaps presented. In panels 3f and g number of simulations is 40 per data point.

## Extended Data

**Extended Data Fig. 1.**
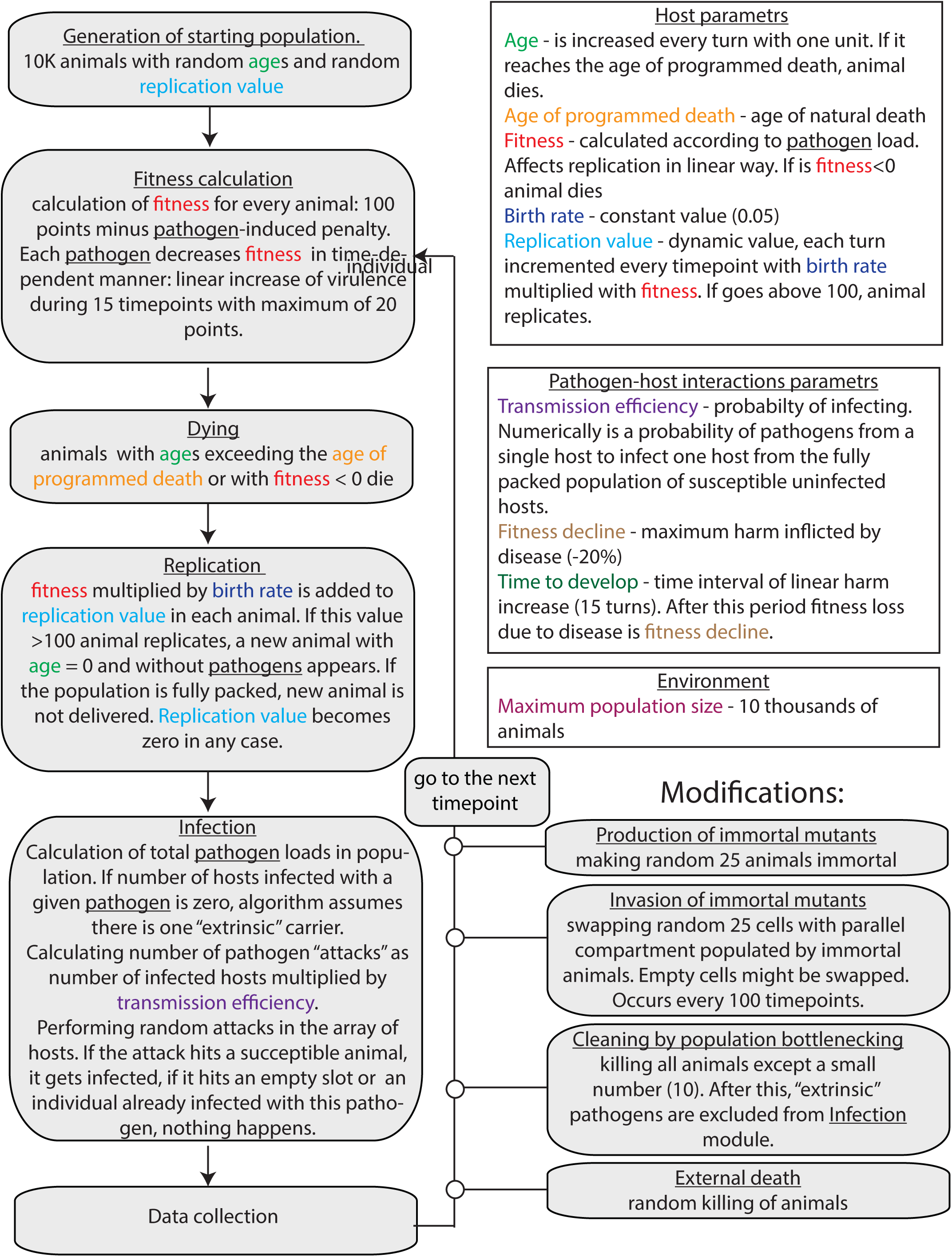
Simulation algorithm.

**Extended Data Fig. 2.**
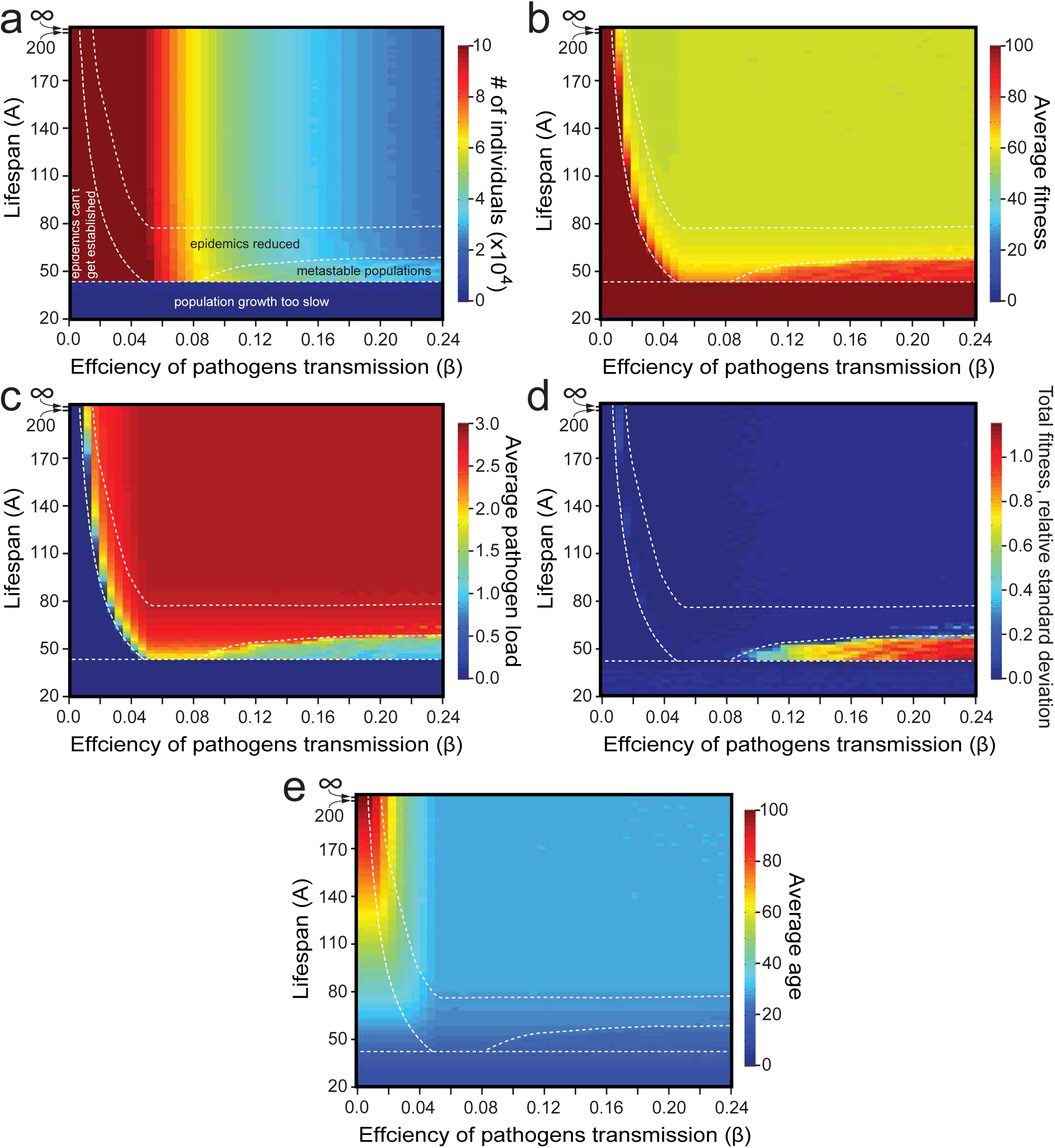
Detailed demographics data of simulated populations from Fig. 1e. **a.** Average number of individuals. **b.** Average fitness. **c.** Average number of pathogens per individual. **d.** Standard relative deviation of total fitness reveals the area of population metastability. See Extended Data Fig 3e-g for more details. **e.** Average age. All data were collected after epidemics had reached a steady state.

**Extended Data Fig. 3.**
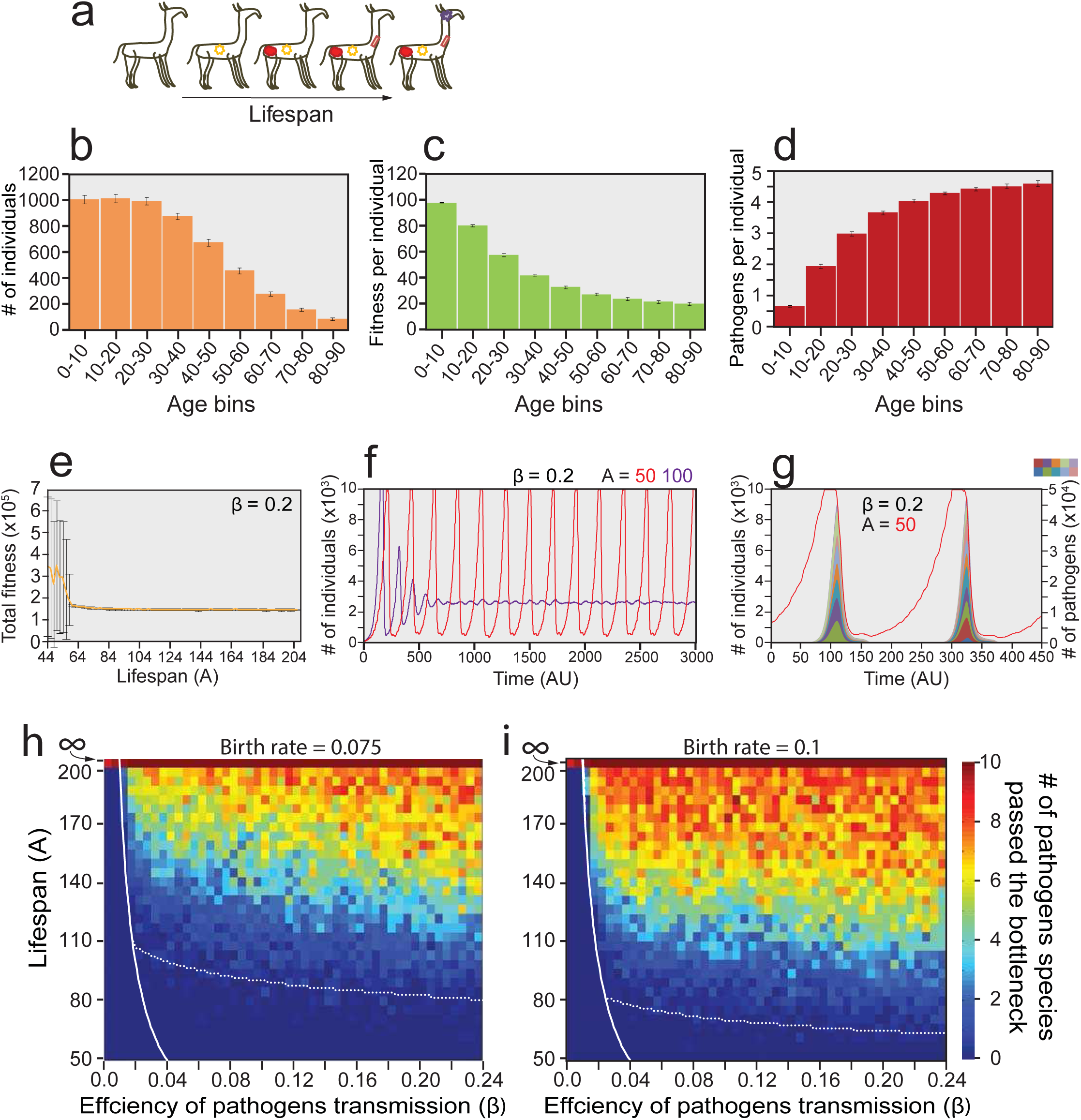
Lifespan setpoints effect on epidemic progression, population metastability and pathogen clearance during host population bottleneck. **a.** A conceptual scheme of lifespan setpoints effects on epidemics control: older individuals have statistically more parasites than their younger counterparts since the probability of getting infected increases with the time. **b-d.** Analysis of individuals stratified by their ages. Simulation was performed with individuals with an age of programmed death of 90 time units, exposed to pathogens with a transmission efficiency of 0.095. Confidence intervals are standard deviations. It should be noted that similar effects were previously reported and discussed as an independent reason for aging evolution^9^. However conservation of lifespan setpoints according to this concept would require severe epidemics to go on constantly in all the species, which is a questionable assumption. **b.** Demographic pyramid of infected populations. **c.** An individual’s average fitness decreases with age. **d.** Average number of pathogens per individual increases with age. Therefore elimination of older individuals increase population fitness. **e-g.** Detailed data on metastable populations. **e.** Total population fitness of individuals with different lifespan setpoints. Standard deviations were high in simulations with short hosts lifespans. **f.** Number of individuals in populations with two different values of lifespan setpoints. **g.** Two peaks from panel f are overlaid with pathogen composition. When the population reaches a certain threshold, epidemics rise, kill a majority of individuals and then vanish. Results are shown for pathogen transmission efficiency of 0.2. **h-i**. Efficiency of pathogen clearance during bottleneck dispersal strongly depends on birth rate. Simulations were carried out as in Fig 2c using an birth rate of 0.075 (**h**) and 0.1 (**i**). Solid lines are from Fig 1e, while dotted lines are predicted borders of pathogen clearance for a given birth rate produced from the deterministic model shown in Fig 2d.

**Extended Data Fig. 4.**
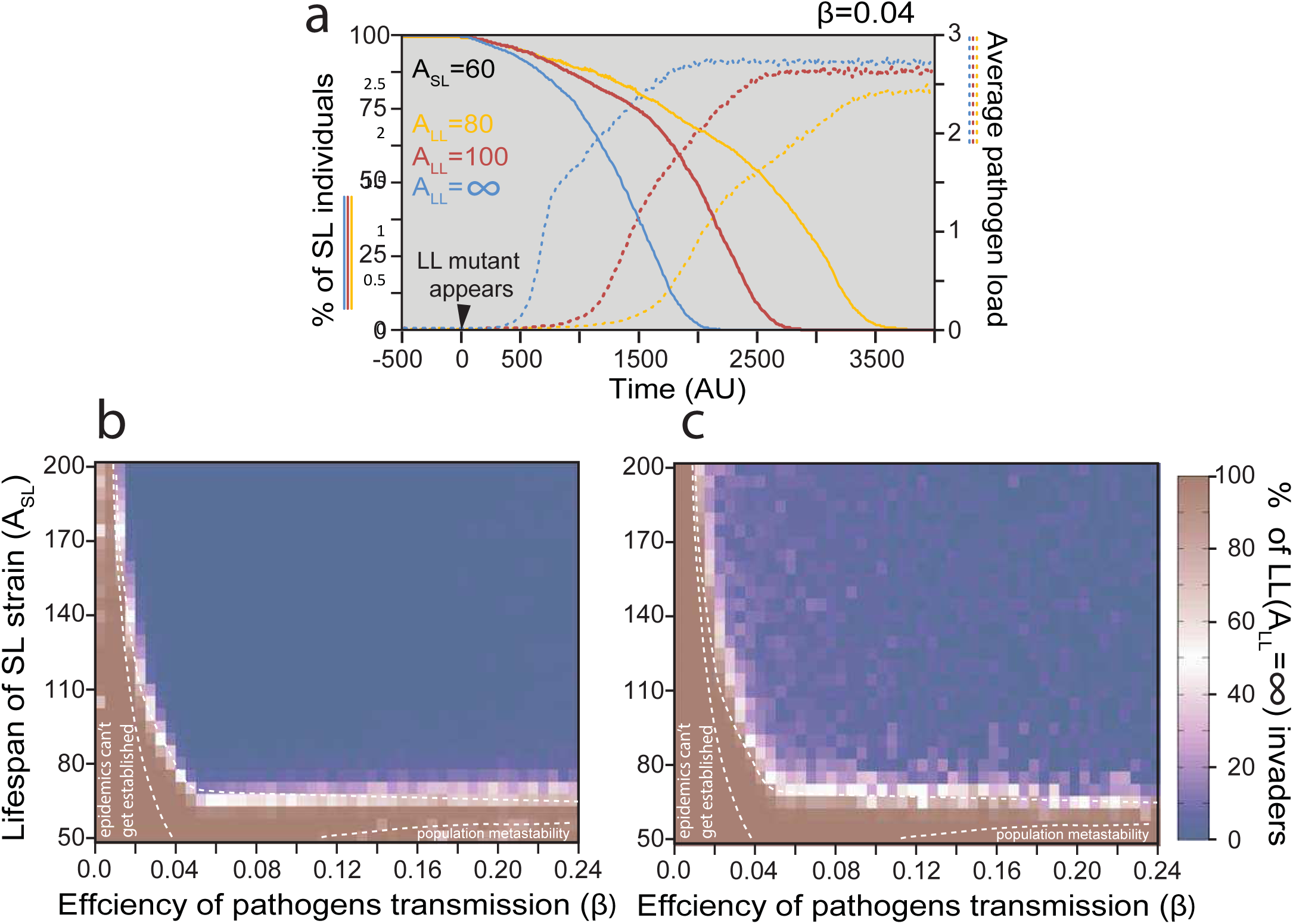
Long-living mutants are efficiently displacing short-living ones in uniformly mixed populations. **a, b.** Populations of individuals with an age of programmed death of 60 were exposed to pathogens with a transmission efficiency of 0.04. At the time point designated by 0, 25 random individuals were transformed into long-living mutants. Dynamics of paternal strain elimination (solid lines) and epidemic explosion (dashed lines) are shown for three different simulations with the longevity of long-living mutants set at 80, 100 or ∞. **c,d.** Systematic analysis of short-living individuals’ displacement by long-living (A=∞) mutants. Percent of non-aging mutants at the last time interval is shown. Dashed lines are from Fig 1e. **c.** *In situ* generation of a long-living mutant. Simulation was performed as in panel *a* with different longevities of short-living mutants and pathogen transmission parameters. **d.** External invasion of a long-living strain. Short-living and long-living populations were run in parallel. After epidemics have reached the steady state, random groups of 25 slots were swapped between two compartments every 100 time points.

**Extended Data Fig. 5.**
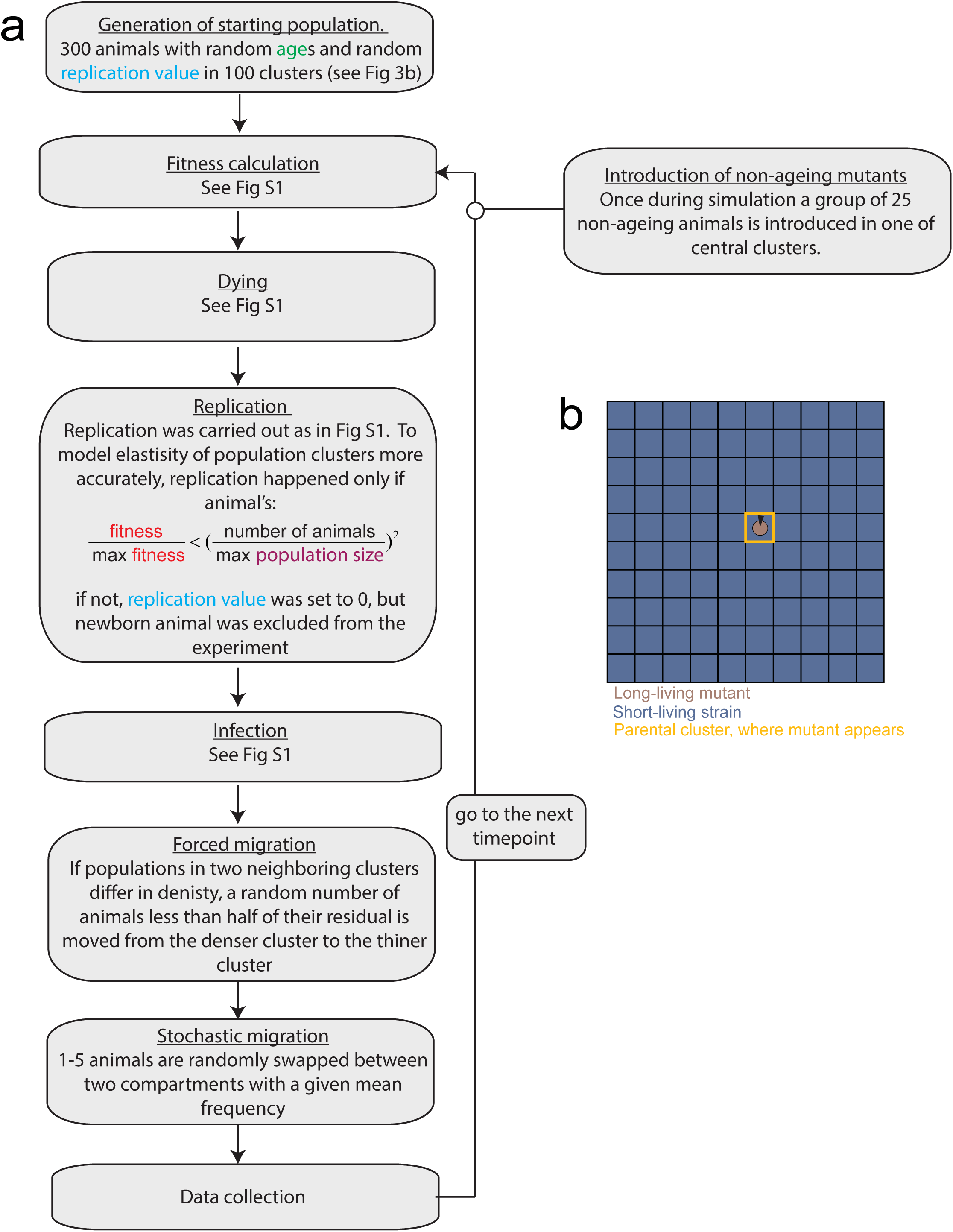
Metapopulation simulations algorithm.

**Extended Data Fig. 6.**
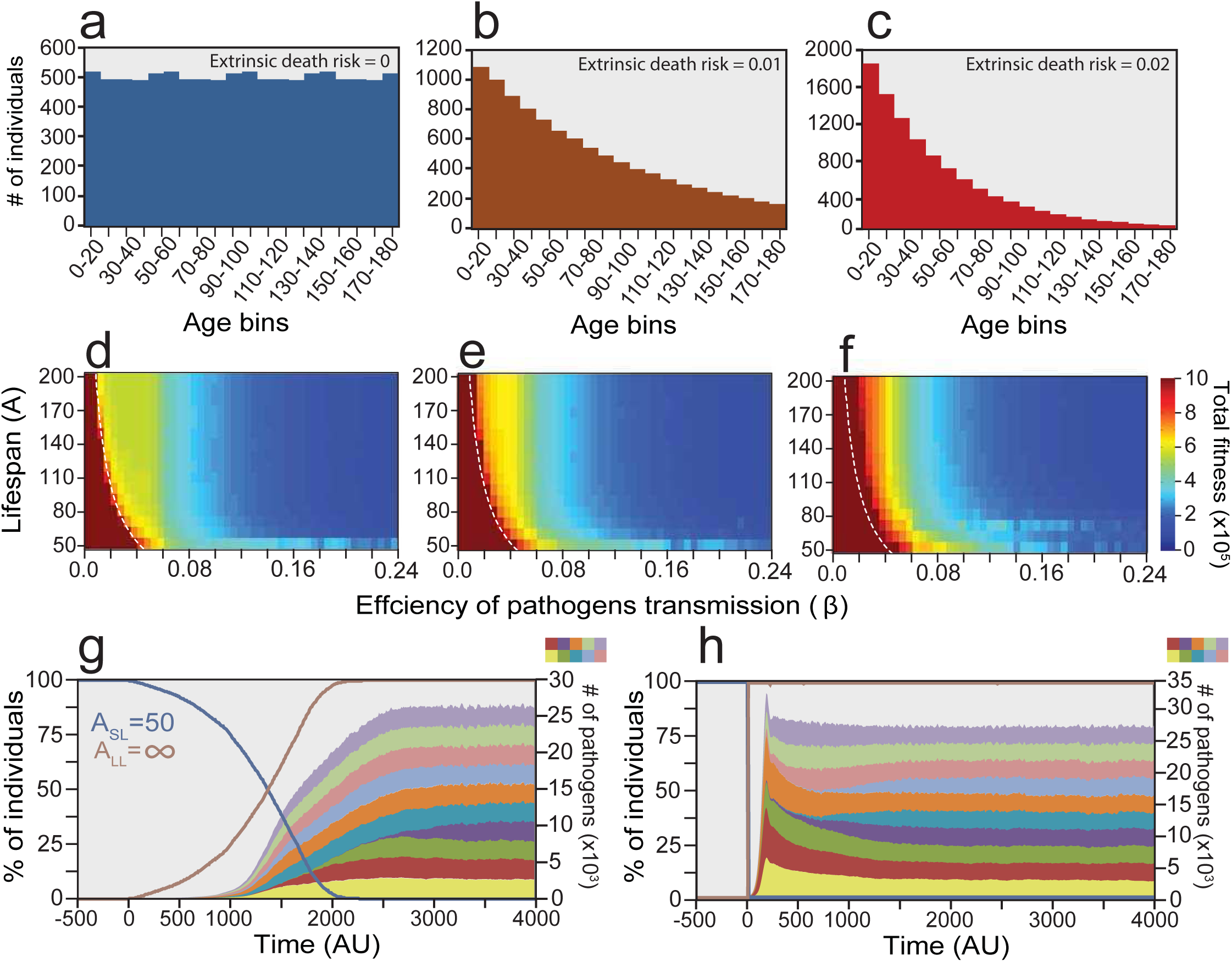
Effects of extrinsic death on epidemics progression. **a-c.** Demographic pyramids of populations exposed to different levels of aging and disease-unrelated death risks. No pathogens were present in these simulations. **d-f.** Populations exposed to high extrinsic death risks are less efficient in establishing epidemics. Simulations were run as in Fig 1e with different values of extrinsic death. Dotted lines are from Fig. 1f. **g and h.** Adverse effects of non-aging mutant production and radical lifespan management. **g.** Production of long-living (A=∞) mutants result in pathogen load increase after 10-15 average lifespans have passed. Displacement of the short-living (A=50) line requires a long period of time. **h.** Transformation of all the short-living individuals into long-living ones (e.g. by removal of extrinsic death or by pharmacological treatment) results in rapid emergence of epidemics, within 2-3 average lifespan lengths. Pathogen transmission was 0.04 in both panels.

**Extended Data Fig. 7.**
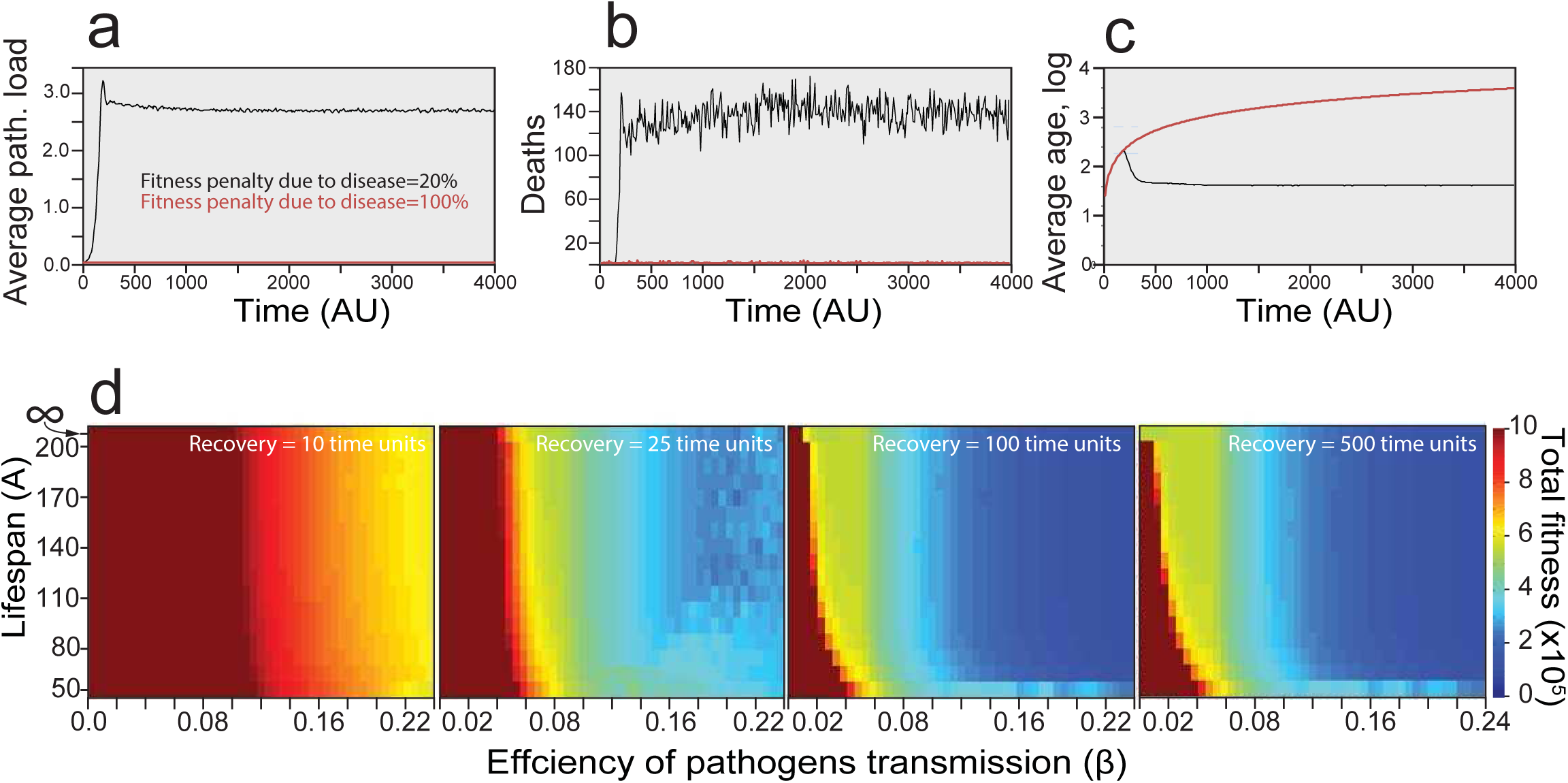
Effects of hypersensitivity and recovery from diseases on aging evolution n. **a-c.** Increase in pathogen-associated fitness penalties prevents epidemics and makes immortality feasible. Simulations of long-living (A=∞) individuals population dynamics were performed with fitness penalties of -20% as elsewhere or with increased (-100%) ones. Average pathogen load (**a**) and death rate (**b**) were clearly minimal in increased fitness penalty situations. **c.** Average individual age was stabilized by disease-inflicted mortality when pathogen-associated fitness penalty was low, but grew throughout the simulation if the penalty was high. Therefore individuals were virtually immortal in the last case. Pathogen transmission efficiency (β) was 0.04 in these simulations. **d.** Aging-related benefits are produced by diseases with slow or no recovery. Simulation experiment as in Fig 1e was performed with diseases having different recovery times. The profile of the region inaccessible for epidemics was convex if the disease was long lasting (recovery = 500 time units, longer than host’s lifespan), and becomes almost vertical (and thus aging-independent) if recovery was fast (10 time units).

**Extended Data Movie. 1. Outcome of competition between short-living and long-living strains in clustered populations depends on the ratio between density-dependent and density-independent dispersal.** Simulations were made with the age of programmed death at 70 time units, pathogen transmission efficiency was 0.03, the probability of stochastic dispersal is shown in the bottom-left corner of each sample. The proportion of long-living (A=∞) individuals in each cluster is shown by color, dark blue corresponds to 0, dark red - to 1.

**Extended Data Table 1.** a. Aging-related observations explained by different evolution theories. Traits which fit the corresponding theory are marked green, traits which contradict the theory are marked red, while traits that are not clearly explained by the theory yet do not appear to clearly contradict it are marked yellow. The prediction itself is marked blue. It should be stated that contradicting facts here are not proving any theory wrong, but rather imply a stronger selection factor that acts against the direction predicted by a given theory.

|  | Damage accumulation | Selection shadow | Disposable soma | Acceleration of evolution | Infectious catastrophe |
| --- | --- | --- | --- | --- | --- |
| Mortality is ubiquitous in advanced animals<sup>a</sup> | Assumes aging as inevitable entropic process | Immortality is absent in species with low external death risks | Immortality is absent from species with slow reproduction rates | Slow evolving species still retain short lifespan | Parasites and infections are ubiquitous factor, affecting all groups of life |
| No immortal or extraordinary long-living animal mutants are observed<sup>b</sup> | Assumes aging as inevitable entropic process | Extreme longevity is impossible even if external death risks are low | Extreme longevity is impossible even if reproduction expenses are low | No strong explanation | Ability to produce an immortal mutant results in elimination of parental strain |
| Extension of lifespan is observed in single-gene mutants<sup>c</sup> | Direct contradiction | Antagonistic pleiotropy-coupling of senescence with beneficial traits | Direct contradiction when fertility is not affected by mutation | aging is adaptive and programmed | aging is adaptive and programmed |
| Hormesis<sup>d</sup> | No strong explanation, no major contradiction | Direct contradiction | Direct contradiction | In a good agreement | In a good agreement |
| aging is observed in wild populationse | In a good agreement | Critical contradiction | In a good agreement | In a good agreement | In a good agreement |
| aging and mortality are absent in some primitive animalsf | Direct contradiction | Some of these animals are having high external death risks | Immortal species do not show decreased fertility | Adaptive aging did not evolved or lost | Adaptive aging did not evolved or lost |
| Metabolism speed and longevity correlate negatively<sup>g</sup> | High metabolic rates result in faster damage accumulation | No strong explanation | Growth of maintenance expenses | Animals with fast metabolism can afford to die faster | Slower metabolism results in a slowdown of low-transmissible pathogens |
| Body size and longevity correlate positively<sup>h</sup> | Larger bodies accumulate more damage, but larger animals are having lower metabolic rates | Big body size protects from predation therefore decreasing external death risks | Big body size is usually associated with low fecundity | Smaller animals can afford shorter lifespan, which is adaptive | Smaller animals can afford shorter lifespan, which is adaptive. Decrease in population density |
| Fecundity and longevity correlate negatively<sup>i</sup> | High fecundity is associated with higher metabolic rate | Compensation for high population losses | Resources, invested in reproduction are the main reason of aging due to underinvestment into maintenance | Adaptive aging accelerates fecundity | Adverse effects in asymmetric dispersal and pathogen cleaning |
| External death risks and longevity correlate negatively<sup>j</sup> | No strong explanation, no major contradiction | High external mortality relaxes natural selection in later ages | No strong explanation, no major contradiction | Direct contradiction | Animals, adapted to lifespan, shortened due to external reasons, evolved faster aging to keep it short if external pressure relaxes |
| Accelerated aging/suicide behavior in semelparous speciesk | No strong explanation | No strong explanation | No strong explanation | Can explain selection for sterility, but not for death | Realization of adaptive aging program |
| Workers/queens longevity differences in social insects<sup>l</sup> | Direct contradiction | Queens are well protected, while workers are not | No strong explanation | Replicating animals live longer and slow down evolution | A few long-living animals are not a problem for infection control |
| Birds longevity<sup>m</sup> | Birds have high metabolic rates | Volation protects from predation | No strong explanation, no major contradiction | No strong explanation, no major contradiction | Volation affects asymmetric dispersal |
| Bats longevity<sup>n</sup> | Bats have high metabolic rates | Volation protects from predation | Low fertility in many bat species | No strong explanation, no major contradiction | Volation, tolerance to infections |
| Naked mole rats longevity<sup>o</sup> | No strong explanation | Queens are protected, workers longevity is atavistic | No strong explanation | No strong explanation | Prediction: hypersensitivity |
| Bowhead whales and greenland sharks longevity<sup>p</sup> | Slow metabolism | Low external death risks | Combination of slow metabolism and high body mass | Combination of slow metabolism and high body mass | Combination of slow metabolism, high body mass and probably low pathogen load |
| Proteus Longevity<sup>q</sup> | Low fecundity | Low external death risks | Low fecundity | Low fecundity | Combination of low fecundity and probably low pathogen load |
| Tuatara longevity<sup>r</sup> | Combination of slow metabolism and low fecundity | Low external death risks | Combination of slow metabolism and low fecundity | Slow metabolism | Combination of slow metabolism, low fecundity and probably low pathogen load |

**b**
| Predictions |
| --- |
| Positive correlation between longevity and mobility<sup>a</sup> |
| Negative correlation between longevity and areal size<sup>b</sup> |
| Negative correlation between longevity and the frequency of contacts between individuals<sup>c</sup> |
| Negative correlation between longevity and exposure to zoonotic pathogens<sup>d</sup> |
| Hypersensitivity to infections in naked mole rats<sup>e</sup> |
| Type I survivorship curves in large flightless ratites<sup>f</sup> |

^a^ Universality of aging defined our historical views on senescence as an unavoidable entropic process alike friction or rust. Theories of **damage accumulation**^10,11^ extend this traditional concept. However, other theories published up to now are not able to explain why virtually all species age. The concept of a **selection shadow**^12^ explains well why aging might be neutral in animals exposed to high external risks of death (e.g. small rodents, insects). However many animals that developed efficient protective mechanisms (e.g. porcupines, skunks etc), large animals (e.g. elephants, rhinos) or predators (e.g. orcas, lions, bears etc) do not have obvious high risks of extrinsic death, but still display senescence. The theory of antagonistic pleiotropy^13^, which usually goes hand-in-hand with selection shadow, proposes multiple senescence-related traits to be genetically linked to early-life beneficial functions, thus resulting in aging conservation. However, such genetic or functional linkages are evolvable properties by themselves, and it remains unclear why evolution cannot overcome this barrier or compensate for the impaired functions by mutations in other regions of the genome. Obvious facts of lifespan evolution, evident from analysis of related species (e.g. mouse deer/baleen whales ~14 / ~100 years respectively; hyraxes/elephants ~12 / ~80 years etc)^14^, make this concept even shakier.

**The disposable soma**^15^ theory proposes that (i) individuals are limited in resources and (ii) faster reproduction rates are favored by natural selection, forcing individuals to accelerate replication and underinvest in maintenance. While being convincing in explaining aging of small exponentially growing organisms, this theory does not exclude full biological immortality. Sufficient amounts of resources might allow maintaining the parent’s body indefinitely if reproductive expenses are low as is true for slow reproducers (e.g. large animals, tuataras etc). We believe it cannot explain aging in high animals which usually have high external death risks in youth, low external death risks in adulthood and stable population numbers. **Acceleration of evolution** theories^16^ cannot explain why slow evolving groups of animals (e.g. sharks or tadpole shrimps) still retain senescence levels comparable to rapidly evolving ones. In our opinion faster evolution is not an adequate benefit for universal aging and mortality taking in account slowness of beneficial mutation accumulation. Furthermore strong selection factors would remove all unfit individuals by itself without the involvement of aging. Strengthening of minor selection factors by costly means of aging will not result in guaranteed long-term success of the species, since these minor factors might turn out to be not very important. In summary, we think that acceleration of evolution is impossible to substantiate outside the framework of altruistic behavior and/or group selection.

**Infectious catastrophe** theory proposes infectious disease as a major factor for the evolution of lifespan setpoints and senescence. Ubiquity of parasites in the biosphere makes aging an almost general property of living matter.

^b^ Lifespan varies dramatically between different species and its evolution is clearly evident across the tree of life^14,17^. On the contrary, intra-species variations are usually moderate. Extremely long-living mutants (~6 fold increase) were described in *C.elegans*^*18*^, however this phenomenon might be attributed to a non-conventional activation of the natural mechanisms underlying roundworm lifespan extension – the so-called dauer stage. Long-living mice mutants display no more than two-fold lifespan extension^18,19^. Being in good agreement with **damage accumulation**, this observation cannot be explained well neither by **selection shadow**, nor by **disposable soma** theories, which consider aging to be a secondary trait, and therefore should be variable across broad ranges. **Acceleration of evolution** is inadequate here as well.

Since longer lifespan produces clear benefits at least within a short evolutionary scale, we should have observed at least some senescence escapers. **Infectious catastrophe** theory is able to explain this by elimination of individuals in the region where long-living mutants emerged. If the fitness penalty associated with immortality or exceptionally long lifespan spreads locally (therefore majorly across the mutant’s parental breed), selection should disfavor even the ability to produce such mutants (Fig 3e).

^c^ Existence of single gene mutants with extended lifespans significantly changed our views on senescence^20–22^. These observations led to scrutiny of the **damage accumulation** group of theories, which assume aging as an inevitable entropic process. **Selection shadow** theory in cooperation with antagonistic pleiotropy could explain these mutant phenotypes since senescence appeared to be linked to stress response and, sometimes, fertility^22,23^. However uncoupling longevity and fecundity directly contradict the **disposable soma** theory. On the other hand, **evolution acceleration** and **infectious catastrophe** theories assume aging to be an adaptive mechanism that obviously can be affected by single gene mutation at least slightly.

^d^ Many environmental conditions can increase lifespan. This includes exposure to heat, oxidative stress, reduction in respiration and translation^22^. The most striking example of hormesis in aging is the effect of caloric restriction on lifespan^24^. Rodents, worms and flies limited in nutrients live significantly longer, demonstrating lower body mass, delayed maturity and sometimes reduced fertility^22^. **Damage accumulation** theories in general do not predict the existence of a pathway for delayed aging. One might speculate, that some specifically harmful metabolic processes slow down under caloric restriction conditions, however this idea does not completely fit the experimental data: dietary restriction in *Drosophila* might prolong life even if applied in late ages^25^. **Selection shadow** theory cannot explain why the existing mechanism of life elongation is not utilized under optimal conditions. Predictions made by the **disposable soma** theory are opposite to experimental observations. Resource deficiency should affect longevity and fecundity unidirectionally in individual life history, e.g. caloric restriction should lead to decrease in lifespan or fertility (or both). Opposite changes are difficult to explain in the frame of this theory. Robin Holliday made an attempt to remove this contradiction by saying that caloric restriction prolongs lifespan by activation of the program for “famine survival”^26^. However it remains unclear why similar programs for aging prevention cannot be activated in *ad libitum* situations when available resources are sufficient for both maintenance and reproduction. To the contrary, **evolution acceleration** and **infectious catastrophe** are in good agreement with tunable aging. If an organism’s death is adaptive, its optimal timing can vary between different environments and change in response to external stimuli. Short-living species reproduction is specifically vulnerable to adverse environmental conditions, e.g. a single poor season might strongly affect the whole species population in case its duration is comparable to the length of the animal’s lifecycle. To survive periods of low productivity, these species could evolve mechanisms to slow down the aging clock in response to hunger, abnormal temperatures or oxidation. In agreement with this idea, signals from the reproductive system also influence aging in some short-living species^22^ – there is probably no ecological benefit to keep animals alive for longer than required for replication. It is important to note that model organisms used as laboratory models for aging research are usually short-living, therefore one should be specifically careful when extrapolating hormesis results on long-living species whose reproduction is less dependent on environmental fluctuations.

^e^ Field studies suggest senescence is detectable in wild animal populations^27^. The effects of aging on mortality could be also extrapolated from the analyses of an animal’s survivorship curves. If the input of aging is minimal, and the majority of individuals die due to extrinsic reasons (e.g. predation, hunger and diseases), the logarithmic curve is linear (Type II, mortality is age-independent) or concave (Type III, mortality is higher in younger ages). If the old age-related reasons play an important role, the curve is convex (Type I, mortality increases with age). Type I survivorship curves are found in many large animal species thus directly supporting that aging plays a role in the wild populations^17^. This goes into a critical contradiction with the **selection shadow** theory, which assumes aging to occur only beyond the period of real survival. All the other discussed theories agree with this observation nicely.

^f^ Several simple metazoans including planarian flatworms (e.g. *Schmidtea mediterranea)*^28^, cnidarian *Hydra vulgaris*^*29*^ and *Turritopsis dohrnii*^*30*^ are considered to be biologically immortal, contradicting all theories of non-adaptive aging. These facts cannot be explained by **damage accumulation** theories, since they show that senescence can be ultimately avoided. A possible argument on existence of exquisite repair mechanisms in these primitive animals immediately raises a question, why have similar adaptations not evolved in more advanced taxa? These species are not protected from external death risks and do not display abnormally low fecundity, therefore challenging both **selection shadow** and **disposable soma** theories. **Evolution acceleration** and **infectious catastrophe** consider that these primordial animals could have lost adaptive aging, perhaps because they did not evolve genetic linkages, preventing this in higher animals. It also might be that aging is mechanistically unavailable for them e.g. due to a functional linkage between regeneration and reproduction.

It is also interesting to speculate about connections between sexual reproduction and aging. Senescence-positive sexual lineages of planarian worms (*Schmidtea mediterranea*) are less efficient in telomere length maintenance than “immortal” asexual ones^28^. We propose that since sexual reproduction requires two animals to contact each other, this opportunity might be used by pathogens for transmission. Therefore asexual species should withstand lower selective pressure towards lifespan shortening.

^g^ Negative correlation between longevity and metabolic rate is widely accepted^31–33^. It is obvious that animals with slow metabolism require a longer time to reproduce. But why do metabolically active animals age faster? This can be easily explained from the positions of **damage accumulation** and **disposable soma** theories, assuming maintenance costs are growing with increase of metabolism rates due to faster accumulation of toxic metabolites, damaged molecules or aberrant mutations. On the contrary, **selective shadow** theory cannot explain this interdependence. **Evolution acceleration** and **infectious catastrophe** theories propose natural selection moves toward adaptive lifespan shortening. Furthermore, infectious catastrophe theory provides an additional level of explanation: since the speed of chronic pathogen reproduction correlates with a host’s metabolic rates, the efficincy of transmission should scale accordingly, relaxing the evolutionary pressure on longevity.

^h^ Correlation between body size and longevity is probably the most obvious life history-related trait^34,35^. Growth and gestation of large animals require prolonged periods of time, but it still remains unclear why smaller animals acquire faster senescence. The positive correlation between size and longevity comes into a contradiction with **damage accumulation** theories, not only because larger bodies should accumulate more errors (e.g.: more cell divisions are required during development, toxic metabolites excretion is more difficult from a bigger volume etc), but also because large animals set the theoretical limits of longevity to a remarkably high level. There is no explanation why damage control systems in small species cannot be as efficient as in bigger ones. A counter argument may address the negative correlation between body size and metabolism levels^36^, however this correlation does not seem to be strong enough to explain huge differences in longevity in different species. For example birds have high metabolic rates and their lifespans are significantly longer than mammals of their sizes (see row ^m^). **Selection shadow** argues that body size better protects from predation, while **disposable soma** theory might suggest a faint argument that body size often negatively correlates with fecundity. In our opinion size/longevity correlation is a strong argument pointing towards shorter lifespan setpoint adaptability: since (i) gigantic species have evolved independently in virtually all the taxa, longer lifespan can emerge without major mechanistic problems, (ii) evolution goes not only towards increases in size, but also vice versa, therefore smaller species that originated from bigger ones should have lost their longevity very efficiently, thus justifying the existence of selective pressure towards lifespan shortening. Thus **evolution acceleration** and **infectious catastrophe** theories are postulating adaptability of aging and therefore explain why animals might limit their lifespan if they physically can replicate within shorter timeframes. In further support of infectious catastrophe theory it should be noted that since increases in body size usually results in decreases of population density^37^, risks of novel pathogen adaptation might go down as well.

^i^ Negative correlation between longevity and fecundity is well established^38^. This observation is the basis for the **disposable soma** theory. However, while longevity/fecundity trade-off seems to be obvious when comparing different species^39^, analysis of individual life histories in some studies did not show great correlation between childbirth and lifespan in mammalian, bird and insect systems^40–43^, suggesting that fecundity is an ecological rather than a physiological factor of aging evolution at least in some cases. **Damage accumulation** theories can explain this with adverse side effects of high energies, individuals invest in reproduction. **Selection shadow** concept might produce an indirect reasoning that high fecundity is compensating for high population losses from external deaths. **Evolution acceleration** and **infectious catastrophe** might explain the longevity/fecundity trade-off on an individual level with a reverse argument of disposable soma theory: unavoidable rapid (adaptive) senescence can be an evolutionary cause for higher fecundity. Indeed the preexistence of lifespan setpoint might force individuals to exhaustively invest in replication as described ^44^. As for as the fitness price of decreasing residual lifespan, its loss can be balanced by one successful act of reproduction. The second argument, specific for short-living species, is signaling from the reproductive system that can manage lifespan (discussed in detail in row ^d^). In addition to this, **infectious catastrophe** theory provides two strong reasons for longevity/fecundity trade-off on ecological level: we claim that high fecundity can (i) degrade short lifespan-related benefits in pathogen clearance (Fig 2g, Extended Data Fig. 3h, i) and (ii) favors shorter lifespan fixation during asymmetric dispersal (Fig 3h to j). Thus we posit a connection between longevity and fecundity as a complex ecological and physiological entity, involving many different factors.

^j^ High external death risk is believed to have a negative correlation with longevity^45^. In this paper we propose revisiting some examples frequently cited as supporting this idea (see e.g. rows ^m, l, o^), but we do not question this rule itself. This correlation is a key element of **selection shadow** theory. **Damage accumulation** and **disposable soma** theories can produce indirect arguments, by linking external death to requirement for high fecundity and therefore high metabolic rates. **Evolution acceleration** theory comes into a direct contradiction with this fact, since the reasons to maintain senescence in the presence of strong selective forces seem to be questionable. **Infectious catastrophe** theory claims that the exact mechanism of an individual’s death does not play a role in pathogen control, and even massive death by predation or hunger might help to cease the epidemics (Extended Data Fig. 6a to f). Animals exposed to high extrinsic death risks adapt their immunity and fecundity to relatively low pathogenic loads as if they had naturally short lifespan. In this situation transitional relaxation of death risks pressure (decrease in carnivore’s numbers or abundant crops) might result in even more severe adverse epidemiological effects than long-living mutant production (Extended Data Fig. 6h). Hence the species exposed to high extrinsic death risks should keep senescence as a reserve security mechanism to kill elderly animals at ages comparable to their death from extrinsic reasons.

^k^ Accelerated aging and suicidal behavior observed in many semelparous species^46,47^ clearly indicate an organism’s death itself (and not just its removal from the next replication round) is favored by selection. All theories except **infectious catastrophe** cannot address this. We consider semelparity is an adaptation of short-living species that guarantee them a successful round of replication and a prompt removal from the population. If their senescence would depend on time only, some individuals would not be able to reproduce due to random variations in the environment. We explain striking examples of accelerated aging in semelparous fish of the *Salmonidae* family^48^ by the necessity to control the spread of parasites in the overcrowded rivers they spawn in. See also row ^d^ for discussion on semelparity.

^l^ Queens of social insects live much longer than workers or soldiers^49^. This fact contradicts the **damage accumulation** and **disposable soma** theories: highly fertile and metabolically active animals are living far longer than sterile individuals from the very same species. The argument of efficient allocation of resources and underinvestment into workers’ maintenance will not work here, since it assumes that building up a new individual from a fertilized egg is cheaper than maintaining an existing one. On the contrary, **selection shadow** explains this by efficient protection of queens and exposure of workers to high external death risks. **Evolution acceleration** theory would expect the replicating animals to have the shortest rather than longest lifespan, coming into a direct contradiction with the observed fact. According to **infectious catastrophe** theory a worker’s senescence is crucial for control of epidemics, while a single long-living queen does not have a major effect due to the low probability of being exposed to a zoonotic pathogen.

^m^ Birds are known to live significantly longer than most mammals of comparable size^34,35^. This comes into a direct contradiction with the **damage accumulation** theories since birds have very active metabolisms. **Selection shadow** theory claims birds are well protected from predation due to their ability to fly and this is one of the most frequent examples mentioned in support of this theory. Large flightless birds have remarkably shorter lifespan^35^, emphasizing the connection between volation and longevity. However, the existence of specialized avivore predators among other birds, reptiles and mammals is well established. Indeed even in classical works describing animals’ life-histories^50^ many species of volation-competent and relatively long-living birds are ascribed to possess Type II survivorship curves, pointing that extrinsic reasons of death play a key role in their mortality. While possessing average lifespancompared to those of small mammals when in the wild, these birds are able to survive for decades in captivity^35^. It is also difficult to consider the songbird’s longevity as atavistic: short-living flightless species are believed to evolve independently from flying ancestors^51^, thus demonstrating an efficient convergent evolution towards lifespan shortening. We consider these arguments to point a contradiction in the selection shadow concept. **Disposable soma** and **evolution acceleration** theories are not capableof producing a strong explanation for birds’ longevity. **Infectious catastrophe** theory predicts that the longevity of birds is associated with their high mobility due to volation. During asymmetric dispersal phases highly mobile animals more readily achievelonger lifespan (Fig. 3a, f, g). Large flightless birds have lost their mobility and therefore experience a stronger pressure towards lifespan shortening. See also Extended Data Table 1b, row^f^.

^n^ Some species of bats live extraordinary long lives for mammals of their sizes^35,52^, even when compared to birds. To the arguments listed in row^m^ we should add that bats have a unique immune system that makes them tolerant to a wide class of pathogens^53^. This fact nicely fits the **infectious catastrophe** theory: tolerance to infections relaxes the pathogen-related pressure towards shorter lifespan.

^o^ Naked mole rats (*Heterocephalus glaber*) are eusocial subterranean rodents of extraordinary long lifespan, which are considered to have negligible senescence or even be biologically immortal^54^.

Moreover mole rat queens remain fertile until the end of their lives. This puzzling observation fits neither damage accumulation, nor disposable soma, nor evolution acceleration theories. The concept of **Selection shadow** might explain it by good protection produced for queens, which reside in the middle of the colony. Workers’ longevity therefore is atavistic. Trying to understand naked mole rats from the **infectious catastrophe** perspective we questioned whether there may be a specific case where biological immortality is evolutionaly stable under non-sterile conditions. We found that an increase in fitness penalties caused by pathogens in our simulations results in their inability to establish epidemics. If the very first individual infected by zoonotic pathogen always dies, the populations are sterile from disease and can evolve towards immortality (Extended Data Fig. 7a-c). Such high penalties cannot be evolutionary stable if the fitness losses are programmed by pathogens, since then infectious agents will evolve towards milder pathogenesis^1^. Fitness loss resulting from a suicidal program of the host in response to infection - hypersensitivity - is clearly an altruistic trait and cannot be fixed by selection.

However, hypersensitivity combined with eusociality might be both beneficial and evolutionary stable. Little is known about naked mole rat’s immunity, however one case report claims these animals are much more sensitive to herpes simplex virus than mice^55^. Our model predicts that this is a pathogen-triggered suicidal phenomenon that should be also true for other infections. We predict that in mole rats the function of pathogen control was shifted from aging to a mechanism of systemic apoptosis (phenoptosis^56^), and thus removing the selection pressure towards shorter lifespan.

^p^ Recent works showed that two huge arctic animals, bowhead whale (*Balaena mysticetus*)^57^ and

Greenland shark (*Somniosus microcephalus*)^58^ have extraordinarily long lifespan of ~200 and ~400 years respectively. These animals are huge and have relatively slow metabolism, therefore their longevity can be explained by all evolutionary theories listed here. From the side of **infectious catastrophe** theory we want to add a probable factor of low pathogenic pressure they experience by living in poorly populated polar seas.

^q^ Proteus (*Proteus anguinus*)^59^ is a long-living (~70 years) troglodyte amphibian inhabiting caves in

southern Europe. Its slow reproduction, absence of natural enemies and life in poor biocoenosis as discussed in row ^p^ makes it easy to explain by all theories.

^r^ Tautaras (*Sphenodon sp.)*^*60*^ are relic New Zealand medium-sized reptiles with an estimated lifespan of

more than 100 years. Their longevity can be attributed to their low fecundity, slow metabolism and absence of natural enemies. It is also tempting to speculate that their biochemical differences from other reptiles, creates additional barrier for pathogens to get established in their populations.

### b. Predictions made by infectious catastrophe theory

^a^ As it is shown in Fig. 3, highly mobile animals are less prone to fix short lifespans when undergoing asymmetric dispersal processes. Thus if all other ecological parameters are the same between two species, less mobile animals should possess shorter lifespans. This prediction is nicely corroborated for an extreme case of mobility – volation. Indeed birds and bats have surprisingly longer lifespans than terrestrial animals of comparable sizes (see Extended Data Table 1a, rows ^m^ and ^n^ for discussion). Further field and laboratory studies are necessary to test this correlation in a broader range of species.

^b^ Range size is also an important parameter in asymmetric dispersal process. Indeed, the same animals inhabiting a small range (e.g. island) will more easily fix longer lifespans since there are not enough refugia, whereas short-lived variants can shelter during initial expansion of a long-living strain. If the long-living mutant conquered the whole range, it will be fixed regardless of the epidemiological consequences. Curiously a comparative study on lifespans of insular and mainland Virginia opossums (*Didelphis virginiana*) directly shows that islanders live significantly longer lives (23% of increase)^61^. However this result cannot be considered as a pure corroboration of the correlation between longevity and range size: the involvement of other factors, such as low pressure from predators (as concluded in^61^, see also Extended Data Table 1a, row ^j^) and parasites (this Table, row ^d^) is not excluded. This prediction has to be accurately tested with multiple different species with control for all of the factors listed above.

^c^ Efficiency of pathogens transmission depends on the frequency of contacts between individuals. Thus if the animals are living as a sparse population with rare contacts between individuals, the model would predict lower probabilities for establishment of new epidemics and therefore longer lifespans. On the other hand, high levels of intermixing would result in shorter lifespans. Solitary, asexual animals are the best candidates to be centenarians. Interestingly, the planarian flatworm considered to be “immortal” is indeed asexual (see Extended Data Table 1a, row ^f^).

^d^ The ability of zoonotic pathogens to adapt to a given species is a key factor of lifespan evolution according to our theory. Therefore frequencies of interspecies contacts are crucial. Thus predation should be an important factor pushing the lifespan towards shortening; however population densities of carnivores are usually smaller than those of herbivores, thereby complicating the analysis. Animals living in abiotic environments such as troglobionts (e.g. proteus, see Extended Data Table 1a, row ^r^) are expected to live longer. One more important factor is biochemical compatibility: highly diverged animals are expected to fix zoonotic pathogens with lesser efficiency. Therefore “living fossils” that do not have many related species in their surroundings should live longer (e.g. tautara, see Extended Data Table 1a, row ^q^). Accurate modeling approaches together with high quality ecological data should support these predictions.

^e^ We predict that naked mole rats are hypersensitive to infections, and this feature is responsible for zoonotic pathogen control making aging redundant (see Extended Data Table 1a, row ^o^). Experimental research should test this hypothesis accurately, although one case report has already shown a similar effect^55^.

^f^ Flightless ratite birds have relatively short lives compared to their flight-competent counterparts. This observation is counterintuitive since these birds are larger in size and have low metabolism even among flightless birds^62^. The classical explanation, based on the selection shadow theory argues that flightless birds are not protected from predators and increased extrinsic mortality leads to the evolution of shorter lifespans, thus implying the survivorship curve in ratites should be of Type II. Our theory explains the ratite’s short lifespans by the loss of mobility associated with volation. Therefore based on our prediction large flightless ratites should have Type I survivorship curves in the wild, as it would be generally expected for animals of their sizes (see also Extended Data Table 1a, row ^m^). Unfortunately high quality field data are not available for these birds. We think that investigations of survivorship curves in wild populations of large flightless ratites, such as common ostriches (*Struthio camelus*), emus (*Dromaius novaehollandiae*), rheas (*Rhea sp.*) and cassowaries (*Casuarius sp.*) are needed to corroborate this prediction.

**Extended Data Table 2.**
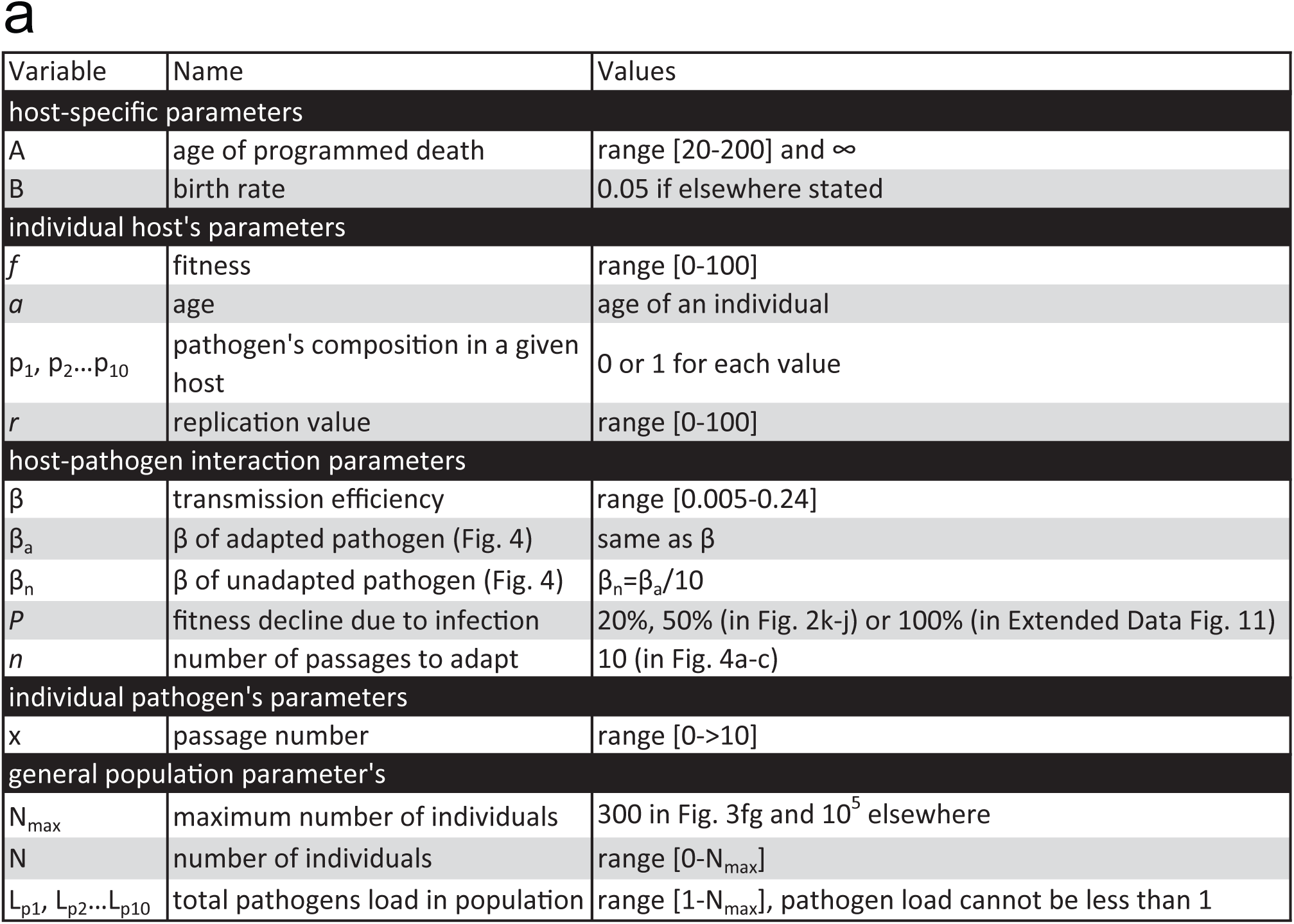
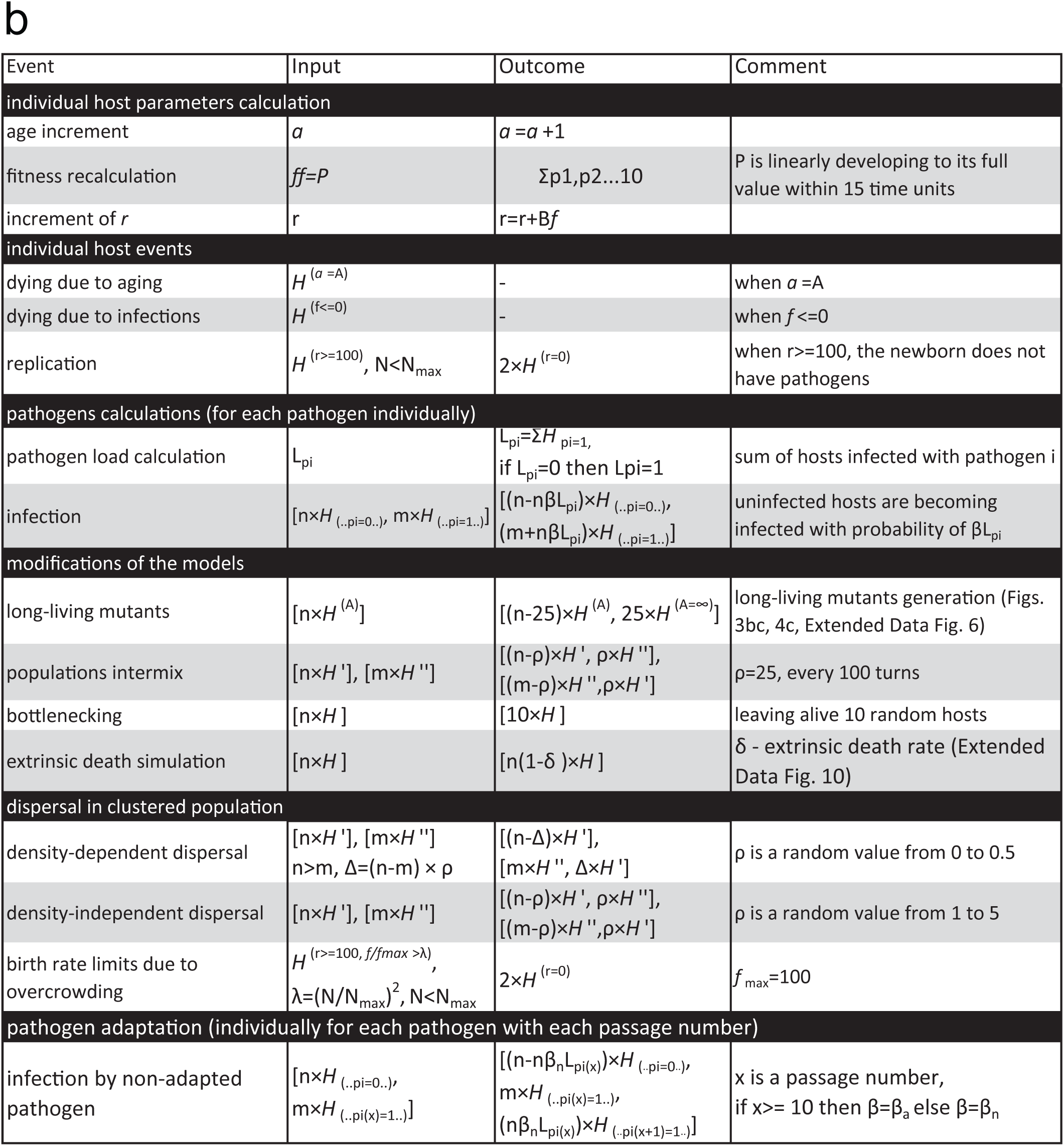
**List of parameters and processes involved in simulations. a.** Parameter’s list. **b.** List of processes. *H* stays for host individual. It’s parameters are shown as superscript, while presence of pathogens in this host, as well as their passage number – as a subscript. E.g. 25×H_(..pi=1..)_ means 25 individuals infected with pathogen *i*. H_(..pi=0..)_ stays for one host not infected with pathogen *i*.

